# Context-specific GITR agonism potentiates anti-PD-L1 and CD40-based immuno-chemotherapy combination in heterogeneous pancreatic tumors

**DOI:** 10.1101/2023.06.16.545301

**Authors:** Chanthirika Ragulan, Krisha Desai, Patrick Varun Lawrence, Yuta Ikami, Mohammed Musheer Aalam, Hari Ps, Nagarajan Kannan, David Cunningham, Naureen Starling, Anguraj Sadanandam

## Abstract

Immunotherapy has shown limited success in pancreatic adenocarcinoma (PDAC) patients. To improve clinical management of cancer, it is crucial to identify alternative immunostimulatory targets associated with mechanisms of tumor evolution to facilitate the development of novel combination immunotherapies. Here we categorized PDACs and other cancers (n>7,500) into subgroups based on immunostimulatory glucocorticoid-induced tumor necrosis factor receptor (TNFR)-related ligand (GITRL) and receptor (GITR) expression: *GITRL*^high^+*GITR*^high^ and *GITRL*^high/low^+*GITR*^low^. We characterized immune evasion mechanisms using immunotherapy preclinical trials in four representative immunocompetent mouse models, finding that the GITR agonist, DTA-1 significantly improved responses in GITRL^high^(+GITR^high^) tumors (n=2). Further characterization revealed increased activation of CD8^+^ T-cells (but not T-regulatory; Tregs cells) and enhanced interferon-γ, immunoproteosome, antigen presentation, and T-cell receptor (TCR) gene expression in DTA-1 responders. *In vivo* clonal tracking using DNA barcoding showed that GITR agonist therapy significantly reduced tumor burden by targeting expansion of heterogeneous PDAC clones and not clone-initiating cells (representing potential resistance). However, emerging GITRL^high^+GITR^high^ epithelial-like oligoclones from the responder model escaped immune surveillance to GITR agonist treatment via increased PD-L1, offering a combined anti-PD-L1, CD40 agonist and DTA-1 immunotherapy regimens (with/without chemotherapy) that further improved responses by decreasing PD-L1^+^ myeloid cells. Conversely, mesenchymal-enriched GITRL^low^ models exhibited primary (intrinsic) resistance to GITR agonist treatment due to reduced T-cells and increased myeloid and/or PD-L1^+^ non-immune cells. These results provide pre-clinical context for GITR+PD-L1+CD40- based personalized immuno-chemotherapy combinations for PDAC.

## Introduction

Immune checkpoint blockade is now an established and effective therapy for many cancer types, but responses are limited in pancreatic ductal adenocarcinoma (PDAC) patients.^1^ This unresponsiveness to immunotherapy partly attributes to intra-tumoral heterogeneity and variable immune-suppressed tumor microenvironment.^2, 3^ However, recent studies have challenged the belief that PDACs are “immune privileged”, and both the innate and adaptive immune responses modulate disease progression.^2, 4–6^ There are therefore opportunities to exploit the immune system for therapeutic gain in PDAC patients.

Immunotherapy aims to shift T-cell homeostasis towards sustained cytotoxic T-cell responses whilst minimizing harmful immune-related adverse events.^7–9^ Nevertheless, patients receiving immunotherapy often experience recurrences due to CD8^+^ T-cell depletion or exhaustion.^10, 11^ Glucocorticoid-induced tumor necrosis factor (TNF) receptor-related protein (GITR/TNFRSF18), a member of the TNF superfamily, is a key stimulatory molecule driving homeostasis in favor of cytotoxic responses. GITR is primarily expressed by resting and activated T-cells, and binding to its ligand (GITRL/TNFSF18) on tumor or antigen-presenting cells (APCs) triggers a co-stimulatory signal that potentiates CD8^+^ effector T-cell function while dampening regulatory T-cell suppression.^12, 13^ Thus, GITR+GITRL represent an important agonist immunotherapeutic target to maintain cytotoxic responses and overcome T-cell exhaustion.^13^

Nevertheless, there are contrasting clinical trial reports related to the response to GITR agonists as monotherapy or in combination with anti-PD1 in melanoma and solid tumors, albeit these different agonists are tolerable in patients.^14–16^ In one of the studies, no response to GITR agonist therapy was observed in immune checkpoint inhibitor pretreated melanoma patients.^14^ These studies suggest a need for more understanding of the context to improve the efficacy of GITR agonist therapy and the best choice and sequence of combination therapy. Besides, the role of GITR agonists in overcoming T-cell exhaustion and immunotherapy resistance caused by immune checkpoint inhibitor therapies other than anti-PD1 and anti-CTLA4 are less studied.^13, 17–19^ Moreover, the mechanism of resistance to GITR agonists in tumors, like PDAC, associated with intra-tumoral (clonal heterogeneity) and immune microenvironmental heterogeneity (context) is expected to help identifying effective combination immunotherapy with GITR agonist. Hence, the goal is to study the intrinsic tumor and microenvironment-specific contexts of GITR agonist response and identify the best combination immunotherapies in PDAC.

Here we stratified human PDACs according to GITR+GITRL expression to understand the potential sub-groups (personalized) for therapeutic exploitation through GITR personalized immunotherapy. To obtain mechanistic insights into GITR signaling, we developed mouse models representing the immune repertoire in human PDACs. Using mouse GITR+GITRL expression to predict responses to GITR agonism, we selected tumors with diverse phenotypic heterogeneities for potential personalized treatment to validate our approach *in vivo* and *in vitro* (cancer-immune cells coculture), ensuring clinical relevance by combining our immunotherapeutic strategy with checkpoint inhibitor (anti-PD-L1), another agonist (anti-CD40) and standard-of-care chemotherapy.

## Results

### GITRL+GITR expression is heterogeneous in PDAC and is associated with immunotherapy response

Firstly, to study immune responses in PDAC *in vivo*, we chose well-known anti-PD-1 therapy and treated a syngeneic mouse model (3275), resulting in no therapeutic response (**Fig. S1a**), like that of patient samples^20^. Hence, we sought to identify immunostimulatory targets that may support T-cell-mediated immune activation in PDAC. We examined four major immunostimulatory receptor-ligand pairs (*GITR*:*GITRL, ICOS:ICOSLG, TNFRSF4 (OX40);TNFSF4 (OX40L), TIGIT:PVR*) in 70 human PDACs from the International Cancer Genome Consortium (ICGC; n=70) transcriptome dataset, of which *GITR*+*GITRL* primarily clustered patients into four distinct immune subgroups (**Fig. 1a-b; Fig. S1b-d**): *GITRL*^high^*GITR^l^*^ow^, *GITRL*^high^*GITR*^high^, *GITRL^l^*^ow^*GITR*^low^, and *GITRL*^low^*GITR*^high^.

**Figure 1.**
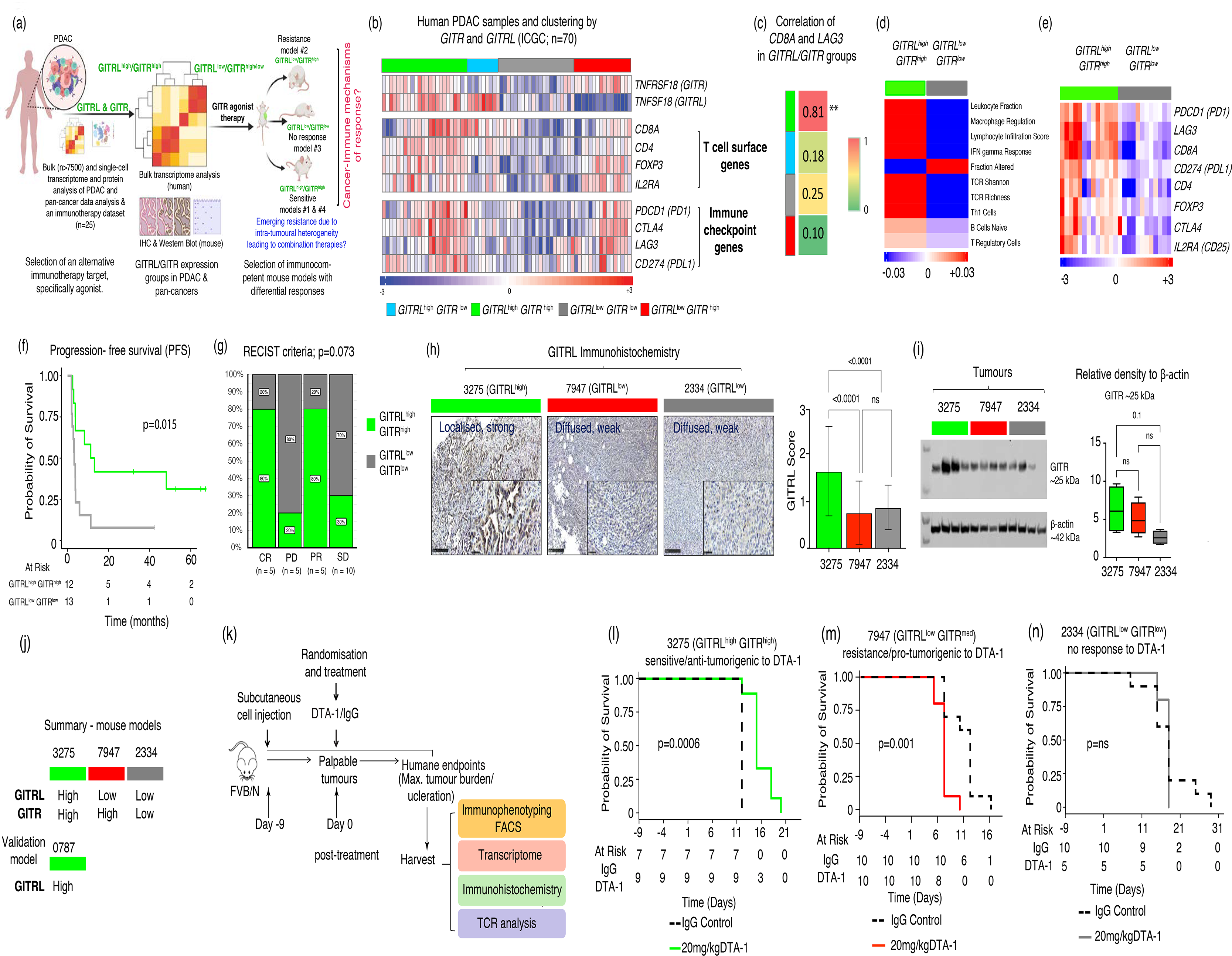
Personalised selection of mouse models for anti-GITR agonist DTA-1 therapy. (a) A schematic showing how to model context-specific differential anti-GITR treatment response in mice based on the classification of PDAC (and pan-cancer) patient samples using GITRL+GITR and T-cell gene expression. (b) A heatmap showing four clusters of patient PDAC (from ICGC) primarily based on *GITRL* and *GITR* high and low gene expression (after median centering). These clusters or groups show differential expression of T cell and immune checkpoint genes. (c) A heatmap of correlation coefficient of CD8A and LAG3 in different *GITRL* + *GITR* groups. ** - represents p < 0.0001 by correlation test. (d) A heatmap showing significant (Benjamini-Hochberg; BH<0.05 by unpaired two-samples Wilcoxson t test) enrichment of various immune cell types and profiles in *GITRL*^high^*GITR*^high^ vs. *GITRL*^low^*GITR*^low^ groups in TCGA patient PDAC samples. Median centered median values of various features across samples and between *GITRL*+*GITR* groups are shown in heatmap. (e) A heatmap showing *GITRL*^high^*GITR*^high^ vs. *GITRL*^low^*GITR*^low^ groups and T-cell and immune checkpoint gene expression (median-centered across samples) in melanoma patient samples treated with adoptive T-cell therapy. (f-g) A Kaplan-Meier survival curve and a bar plot showing f) prognosis (progression-free survival; log-rank test) and g) response to therapy (by RECIST criteria; Chi-square test was used for statistical test) in melanoma patient samples treated with adoptive T-cell therapy. (h-i) Expression and quantitation of mouse GITRL and GITR by h) IHC and i) Western blot in syngeneic mouse models of PDAC. Multiple comparisons of ordinary one-way Anova was performed for statistical analysis. (j) Schematic showing the summary of expression of GITRL and GITR in three mouse models. (k) Schematic representing the preclinical trials with GITR agonist treatment and endpoints. (l-n) Kaplan-Meier survival plots (log-rank test was used for p-value) showing survival of mice from l) 3275, m) 7947 and n) 2334 PDAC models treated with GITR agonist as monotherapy.

GITRL and GITR were primarily expressed on myeloid cells and activated CD8^+^/CD4^+^ T cells, respectively, as assessed in bulk and single-cell data from mouse and human PDACs and breast tumors (**Fig. S1e and S2a-d**). Indeed, the *GITRL*^high^*GITR*^high^ subgroup was enriched for T-cell markers (*CD8A, CD4, FOXP3*, and *CD25* (*IL2RA)*), immune checkpoint and exhaustion genes (*PDL1 (CD274), LAG3, PD1 (PDCD1)*), and *CTLA4* (**Fig.1b; Fig. S3a-g**). These features were further validated in additional PDAC (n=525) and pan-cancer transcriptomic datasets (n>9,000; comprising 32 cancer types, including PDAC), suggesting similar phenomenon across cancers (**Fig. S3i-j**). *CD8A* expression highly correlated with the exhaustion gene, *LAG3* in the *GITRL*^high^*GITR*^high^ subgroup in PDAC (**Fig 1c**; p < 0.05). Concordantly, CD8^+^ T-cells with high *LAG3* expression (and proliferative T-cells with *MKI67* gene expression) showed increased *GITR* expression in single-cell data in breast cancer (**Fig. S2b**). These results suggest that the GITRL^high^GITR^high^ subgroup may be enriched for exhausted CD8^+^ T cells in different cancer types.

We next assessed which immune cell types and processes were associated with the *GITRL*^high^*GITR*^high^ subgroup using TCGA PDAC transcriptome data (n=133). The *GITRL*^high^*GITR*^high^ sub-group showed significant (Benjamini-Hochberg-adjusted p-value; q<0.2) enrichment of leukocyte/lymphocyte and tumor-infiltrating regional fraction and their associated immune responses of increased T-helper (Th)1, cytolytic score (CTA), interferon-gamma (IFNG), T-cell receptor (TCR) richness, and Shannon diversity and B-cells and T-regulatory cells compared with other subgroups (**Fig. 1d and Fig. S3h**). These data further suggest that high GITRL+GITR expression is associated with increased T-cell activation in PDAC.

GITR agonists have been shown to boost cytotoxic CD8^+^ T-cell activities in an adoptive T-cell therapy model of murine fibrosarcoma.^21^ We therefore evaluated the implications of constitutive *GITRL* and *GITR* expression in metastatic melanoma patients (GSE100797; cross-cancer analysis as no such PDAC dataset is available) treated with adoptive T-cell therapy using gene expression data.^22^ Interestingly, T-cell surface and immune checkpoint genes were upregulated and progression-free survival (PFS; p<0.02) and overall survival (OS; p<0.1) were longer in *GITRL^high^GITR^high^*(n=12) patients compared with those belonging to *GITRL^low^GITR^low^*subgroup (n=13; **Fig 1e-f**; **Fig. S3k-l**). Moreover, 80% (8/10) of *GITRL^high^GITR^high^*patients showed complete/partial clinical responses as assessed by Response Evaluation Criteria by Solid Tumors (RECIST)^23^ criteria compared with 33% (5/15) *GITRL^low^GITR^low^* patients showing progressive and stable disease (**Fig. 1g**). In addition, *GITRL^high^GITR^high^* patients showed modest (p=0.15) reductions in tumor regression compared with *GITRL^low^GITR^low^* patients (**Fig. S3m**). These observations with a noteworthy, but a possible small cohort of samples, suggest that quantifying GITRL+GITR may provide an opportunity to stratify patients according to responses to immunotherapy, especially therapies that boost T-cells (like agonistic therapy).

### Syngeneic mouse models show context specific GITR agonist therapy response

Initially, we tested the activity of an anti-mouse monoclonal agonist of mouse GITR (DTA-1) in the well-known PDAC genetically engineered mouse model (GEMM) - *Pdx-Cre; LSL-Kras^G12D/+^; LSL-p53^R172H/+^*(KPC) mice. We observed a heterogeneous and only a borderline significant improvement in survival of mice treated with DTA-1 compared to its saline control treatment (**Fig. S3n**). This suggests, like in patient samples,^14–16^ that DTA-1 has some effect in PDAC, however, it is likely that its benefit depends on certain contexts, potentially intra-tumoral heterogeneity and increased GITRL+GITR expression.

To test this hypothesis driven by the clinical^14–16^ and GEMM data, we next tested whether GITRL+GITR expression is useful for selecting syngeneic PDAC mouse models for anti-GITR agonist therapy. We first assessed expression of GITRL and GITR proteins in three syngeneic PDAC transplanted mouse models and validated in fourth model (immunocompetent; using 3275, 7947, 2334, 0787 cell lines derived from GEMM - *p48-Cre; LSL-Kras^G12D/+^; LSL-p53^R172H/+^* (KPC-p48) or *p48-Cre; LSL-Kras ^G12D/+^; Ink4a/Arf ^flox/flox^* (KIC) mice. Immunohistochemistry (IHC) analysis showed that GITRL+GITR were strongly (GITRL; p<0.0001 compared to other models) expressed in 3275 tumors, so this model was defined as the GITRL^high^GITR^high^ group. By contrast, 7947 and 2334 tumors showed diffuse, weak expression of GITRL but high and low GITR expression, respectively, so these mice were defined as GITRL^low^GITR^med^ and GITRL^low^GITR^low^, respectively (**Fig. 1h-i; Fig. S3o**).

We next tested the efficacy of DTA-1 *in vitro* and *in vivo*. There was significant upregulation of CD8^+^ (%CD45^+^ cells) T-cells in activated mouse splenocytes (IL-2/concanavalin-A-activated; see Methods) exposed to 3275 organoid-conditioned media treated with DTA-1 compared with the immunoglobulin (IgG) control as measured by flow cytometry (**Fig S4; Fig. S5a-c**). These data support the hypothesis that the GITR agonist (DTA-1) modulates T cells *in vitro* in 3275 model.

We next assessed efficacy of DTA-1 *in vivo* in the three syngeneic subcutaneous tumor mouse models expressing different GITRL+GITR levels as a dose response study, with mouse survival and tumor volume changes used as endpoints (**Fig. 1j-n; Fig. S5d-f**). Significantly, responses to DTA-1 (compared with IgG control) were different in the three models. GITRL^high^GITR^high^ 3275 mice lived longer (log rank, p=0.0006) (**Fig. 1l**) than GITRL^low^GITR^low^ 2334 mice (log rank, p value not significant; **Fig. 1n**) treated with DTA-1.

Furthermore, we tested a fourth mouse model (0787 orthotopic; **Fig S5g**) with GITRL^high^, which showed similar response to DTA-1 compared to controls as 3275 model. Contrarily, GITRL^low^GITR^med^ 7947 mice (log rank, p=0.001) (**Fig. 1m**) lived shorter with DTA-1 compared with control treatment. These results suggest that PDAC tumors show differential responses (including a detrimental effect) to DTA-1 treatment and that this response may be determined by GITRL+GITR expression.

#### DTA-1 treatment enhances effector T-cell function but not Tregs in GITRL^high^GITR^high^ PDACs

In addition to the survival difference (**Fig. 1l**), there was a reduction in tumor volume in response to DTA-1 therapy in the 3275 subcutaneous model compared with IgG controls. This was assessed by both luminescence imaging (p<0.05) and caliper measurements (p=0.1; **Fig. 2a-b**). In an orthotopic and interventional 3275 model, there was a modest, with borderline significance (p=0.2), reduction in tumor weight in DTA-1-treated 3275 mice compared with controls (**Fig. S6a-b**).

**Figure 2.**
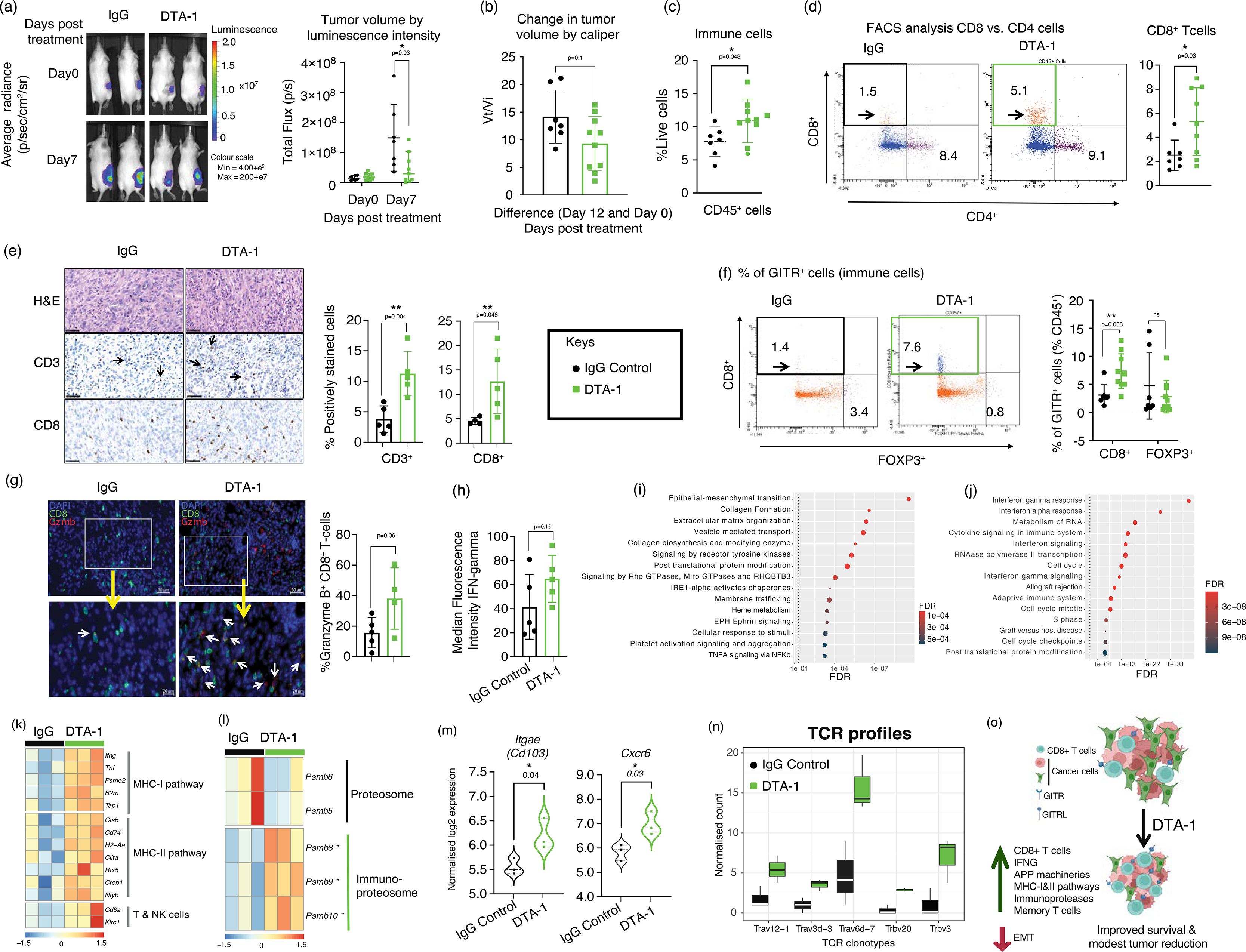
Change in tumor microenvironment and mechanism of response to anti-GITR agonist DTA-1 therapy in 3275 mouse model. (a) Representative figure and quantitation of tumor (subcutaneous) volume as luminescence intensity in mouse tumors treated with IgG or DTA-1. Luminescence was measured using IVIS® Lumina X5 system. (b) Change in subcutaneous tumor volume measured manually by caliper in 3275 mice treated with IgG or anti-GITR agonist therapy. Vi – tumour volume initially and Vt – tumour volume at day 12 after treatment. (c-d) Flow cytometry-based quantitation of c) CD45^+^ immune and d) CD4^+^ and CD8^+^ T cells in 3275 mice treated with IgG or DTA-1. (e) IHC (representative figure)- based quantitation of CD3^+^ and CD8^+^ T cells in 3275 mice treated with IgG or DTA-1. (f) Flow cytometry-based quantitation of CD45^+^ GITR^+^ CD8^+^ (cytoxic T-cells) or CD45^+^ GITR^+^ FOXP3^+^ (T regulatory cells) in 3275 mice treated with IgG or DTA-1. (g) Immunofluorescence and dual-staining quantitating granzyme B^+^ CD8^+^ T-cells in 3275 mice treated with IgG or DTA-1. (h) Luminex results with quantitation of IFNγ in 3275 mice treated with IgG or DTA-1. (i-j) Differential enrichment (hypergeometric; using hypeR package) analysis of gene sets and pathways in 3275 mice treated with i) IgG or j) DTA-1. FDR p value was reported. (k-l) Heatmaps showing differential expression of k) MHC-I, −II and T/NK cells and l) proteosomes, including immunoproteosomes (selected after SAM analysis) in DTA-1 treated and control tumors. (m) Violin plots showing expression of T resident memory cell genes, *Itgae* (*Cd103*) and *Cxcr6*. (n) Absolute count of TCRα/β clonotypes in IgG or DTA-1. (o) Schematic summarizing microenvironment and cancer cell changes associated with DTA-1 treatment in GITRL^high^GITR^high^ mouse PDAC. Unpaired t test was performed for p values for figure a-h and m.

To better understand DTA-1-mediated T-cell responses, we next assessed CD45^+^, CD3^+^, and CD8^+^ T-cells by flow cytometry and IHC, which were significantly (p<0.05) upregulated on treatment (**Fig. 2c-e**; and by gene expression; **Fig. S6c**). There was also a DTA-1-mediated increase in GITR^+^CD8^+^ T-cells but no change in GITR^+^FOXP3^+^ and to lesser extend in CD4^+^ T-cells (**Fig. 2f and S6d**). The activation of CD8^+^ T-cells is evident with modest, with borderline significance, increase in cytotoxic granzyme-B (**Fig. 2g**) and IFN-γ levels (**Fig. 2h**). This suggests that the GITR agonist increases cytotoxic GITR^+^CD8^+^ T-cell infiltration into tumors.

#### DTA-1 remodels tumors towards epithelial characteristics with increased antigen presentation and processing

In cancer cells, DTA-1 treatment also upregulated the epithelial differentiation marker, *Tff1* (with borderline significance; **Fig. S6e**). Hence, we performed pathway analysis of genes differentially expressed (by Systematic Analysis of Microarrays; SAM) between IgG control and DTA-1-treated samples to further understand change in molecular mechanisms associated with DTA-1 treatment. Control samples showed increased epithelial-mesenchymal transition (EMT), extracellular matrix proteins, receptor tyrosine kinases, ephrin, and NFκB signaling (**Fig. 2i**), suggesting that DTA-1 reduces EMT in the GITR^high^GITR^high^ model. By contrast, there was significant enrichment of interferon alpha and gamma signaling, adaptive immune cells, and cytokine signaling (MSigDB hallmarks, KEGG, Biocarta and REACTOME pathways; **Fig. 2j**). Increased effector function is preceded by antigen processing and major histocompatibility complex (MHC)-I and −II presentation (APP).^24^ Accordingly, APP was also enriched in DTA-1-treated samples (but not controls) according to gene set enrichment analysis (GSEA) and KEGG pathway analysis, with many of the included genes involved in MHC-I, −II, and CD8 activation pathways (**Fig. 2k; Fig. S6f-h**). In addition, there was a switch in constitutive proteosomal genes (*Psmb5* and *Psmb6*) to immunoproteosomal genes (*Psmb8*, *Psmb9*, *Psmb10*) in DTA-1 treated samples compared with control mice (**Fig. 2l**). Moreover, there was increased expression of tissue-resident memory T-cell genes (*Cd103* (*Itgae*) and *Cxcr6;* **Fig. 2m**) and five TCRα/μ genes in DTA-1 treated mice vs. controls (**Fig. 2n**). Overall, **Fig. 2o** summarizes DTA-1-specific events leading to anti-tumor immunity in this model. DTA-1 modulated the pro-inflammatory and anti-tumor adaptive immune system in the GITRL^high^GITR^high^ 3275 PDAC model.

#### Intra-tumoral heterogeneity imparts emerging and persistent oligoclones resistant to anti-GITR therapy

The transcriptomic profiling of 3275 tumors treated with a single dose of DTA-1 revealed a significant increase in the expression of *Lag3*, a marker associated with T-cell exhaustion (**Fig. S6i**). This finding suggests the potential development of acquired resistance to the drug. To further investigate this resistance, we administered multiple doses of DTA-1 in the 3275 model. However, we observed only modest improvements in survival benefits and tumor volume changes between the treated and control groups, indicating the evolution of the model towards resistance (**Fig. S6j-k**).

In relation to this acquired resistance, the untreated 3275 tumors exhibited heterogeneity, with adjacent regions within the same tumor displaying varying degrees of differentiation (**Fig. S7a**). To systematically assess this heterogeneity and track the evolution of clones, we utilized our DNA barcoding technology (**Fig. S7c)**. We orthotopically injected barcoded 3275 cancer cells into mouse pancreases and treated them with either DTA-1 or a control vehicle (**Fig. 3a**). There was a significant (p<0.05) reduction in estimated barcoded cancer cells per tumor and clone size in DTA-1-treated mice compared with controls (**Fig. 3b-c**). Consistently, there was a significant (p<0.05) reduction in CD45-negative cells that included cancer cells (**Fig. S7b**). Nevertheless, there was no significant change in tumor clone-initiating cells, suggesting that they are not targeted by DTA-1 (**Fig. 3d**). This DNA barcoding and flow cytometry approaches allowed us to study the behavior of the barcoded cells and gain insights into the effects of DTA- 1 treatment on the evolving clones. These results suggest that 3275 tumors are heterogeneous, and DTA-1 affects cancer cells but not clone-initiating cells, potentially supporting the development of therapy resistance.

**Figure 3.**
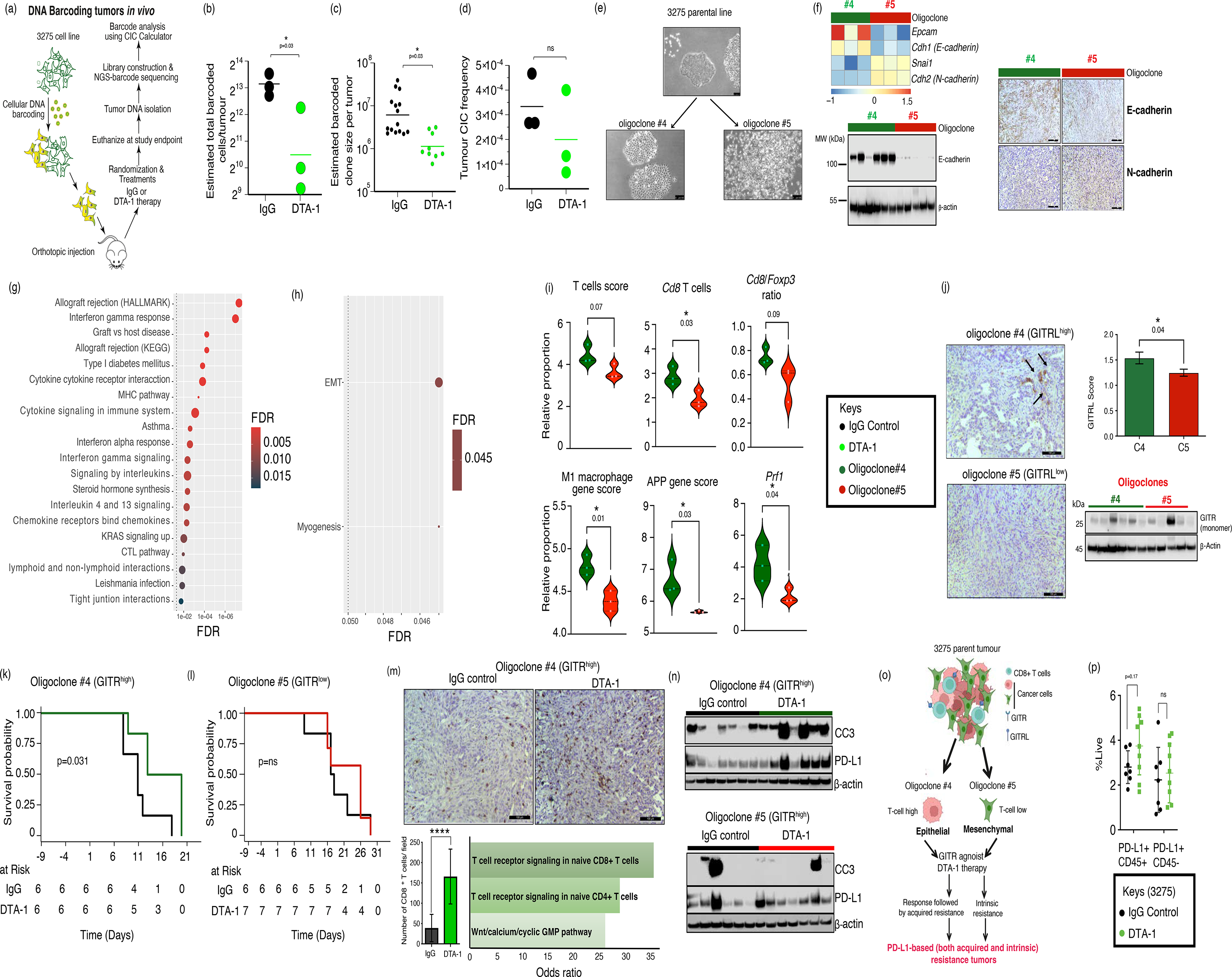
Intra-tumoural heterogeneity associated with resistance to anti-GITR agonist DTA-1 therapy in 3275 model. (a) A schematic representing DNA barcoding analysis in 3275 orthotopic model treated with IgG or DTA-1. (b-d) DNA barcoded results representing estimated total barcoded b) cells/tumour of tissue, c) clone size per tumour, and d) total clone-initiating cell (CIC) frequency. (e) Phase-contrast microscopy pictures showing intra-tumoural cell line heterogeneity showing morphologically different types of cells from 3275 parental cell line. Oligoclones#4 and #5 picked from the parental line with distinct morphologies. Magnification is 100μm. (f) Heatmap, Western blot and IHC showing differential expression of epithelial and mesenchymal markers in oligoclones at gene and protein levels. (g-h) Dot plots showing enrichment (hypergeometric test) of gene sets and pathways in g) oligoclone #4 and h) oligoclone #5 mouse tumours. (i) Violin plots showing differential T-cells, macrophages and MHC-I & −II genes in oligoclone#4 and oligoclone#5 mouse tumours (represented as mean expression scores). (j) IHC for GITRL and Western blot for GITR protein in oligoclone#4 and oligoclone#5 mouse tumours are shown. (k-l) Kaplan-Meier curves (log-rank test for p value) showing differential response of oligoclone#4 and oligoclone#5 mouse tumours to IgG and DTA-1. (m) IHC and gene enrichment showing expression of CD8^+^ T cells oligoclone#4 mouse tumours treated with IgG or DTA-1. (n) Western blots showing differential expression of cleaved caspase-3 and PD-L1 proteins in IgG control and DTA-1 in oligoclone#4 and oligoclone#5 mouse tumours. (o) Schematic summarizing selecting of oligoclones #4 and #5 from 3275 cell lines and their characteristics, including cancer and tumour microenvironment, and response/resistance to DTA-1 treatment. (p) A plot showing change in CD45^+^PDL1^+^ and CD45^-^PDL1^+^ population in 3275 parent treated with IgG control and DTA-1. Unpaired t test was done for b, c, d, i, j, m and p.

To gain a deeper understanding of heterogeneity and resistance to DTA-1, we conducted an *in vitro* selection process to isolate three distinct oligoclones (#1, #4, and #5) from the parent 3275 cell line. These oligoclones exhibited different morphological characteristics, as depicted in **Fig. 3e** and **Fig. S7d**. Oligoclones #1 and #4 displayed an epithelial-like phenotype. Oligoclone #4, specifically, was chosen as a representative epithelial oligoclone for further analysis. On the other hand, oligoclone #5 exhibited a mesenchymal-like phenotype, although it did not possess the typical characteristics of mesenchymal cells. When injected subcutaneously in immunocompetent mice, epithelial markers *Epcam* and *E-cadherin* were variably but significantly increased in oligoclone #4 tumors, while oligoclone #5 tumors showed increased and variable *Snai1* and *N-cadherin* mesenchymal marker expression (by gene and protein levels; **Fig. 3f; Fig. S7e-g**). Interestingly, anti-tumor immunity was significantly increased in oligoclone #4 tumors, while oligoclone #5 tumors showed increased EMT (**Fig. 3g-h**). Correspondingly, gene scores associated with CD8^+^ T-cells, M1 macrophages, MHC-I, and MHC-II were increased in oligoclone #4 tumors compared with oligoclone #5 tumors (**Fig. 3i**), and there was enrichment of gene expression representing adaptive immune responses and tight junction interactions in oligoclone #4 and mesenchymal characteristics in oligoclone #5 tumors (**Fig. 3g-h**).

GITRL expression was significantly increased in oligoclone #4 tumors compared with oligoclone #5 tumors *in vivo*, although they both expressed GITR (**Fig. 3j**). Hence, we examined if oligoclone #4 and #5 contribute to emergent and persistent clones, respectively, to impair responses to DTA-1 treatment in parental 3275 mouse tumors. First, we performed direct co-culture of each oligoclone and activated splenocytes and treated them with DTA-1 or vehicle (IgG; **Fig. S7h**; parental cell line used as a control). We removed splenocyte in suspension to recover the cancer cells after co-culture and treatment. There was almost no change in cleaved caspase-3 (an apoptosis marker) in treated and control oligoclone #5 cancer cells. By contrast, DTA-1 treatment increased cleaved caspase-3 in oligoclones #1 and #4 (**Fig. S7i)**. This suggests that oligoclone #5 may represent a persistent clone resistant to DTA-1- induced apoptosis.

To further test emergence and persistence of the oligoclones in response to DTA-1 treatment, we injected oligoclones #4 or #5 subcutaneously into immunocompetent mice. Oligoclone #4 but not oligoclone #5 mice treated with DTA-1 lived significantly (p<0.05) longer than controls (**Fig. 3k-l**), which attributed to increased CD8^+^ T cells infiltrating this oligoclone (**Fig. 3m**). Again, immunoblotting showed increased cleaved caspase-3 in oligoclone #4 tumors but not oligoclone #5 tumors treated with DTA-1 compared with controls (**Fig. 3n**). Overall, these results suggest that epithelial-like oligoclone #4 was relatively sensitive to DTA-1 therapy whereas mesenchymal-like oligoclone #5 showed persistent resistance to DTA-1 (**Fig. 3p**).

#### Resistance to DTA-1 occurs via PD-L1 expression offering combined anti-PD-L1 therapy in GITRL^high^GITR^high^ PDAC model for improved response

To explore the mechanism underpinning resistance in oligoclones, we assessed PD-L1 protein expression in tumors *in vivo* and cancer cells *in vitro* from direct co-culture of 3275 and splenocytes (**Fig. 3n** and **Fig. S7h, j-l)**. Interestingly, PD-L1 expression modestly increased in DTA-1 treated oligoclone #4 tumors and cancer cells compared to their controls but not in oligoclone #5 tumors, which showed a constitutive expression of PD-L1 (**Fig. 3n and S7j-l**). Indeed, treatment of cancer-splenocyte direct co-cultures *in vitro* with anti-PD-L1 showed modest increase in cleaved caspase-3 expression in oligoclones #4 and #5 and parent compared with control (**Fig. S7m)**. This suggests that secondary (acquired) resistance in oligoclone #4 as an emergent clone and primary (intrinsic) resistance in oligoclone #5 tumors as a persistent clone may be via increased PD-L1 expression in the tumor microenvironment that cumulatively increases in parental 3275 tumors after DTA-1 treatment. Resistance via PD-L1 overexpression may therefore be overcome through combined anti-PD-L1 and GITR agonist treatment (**Fig. 3o**).

Since both tumor and immune cells express PD-L1, we parsed the cell populations based on CD45 into immune and non-immune cells to check the PD-L1 expression in these cell types. DTA-1 modestly (p=0.17) increased CD45^+^PD-L1^+^ (immune cells with PD-L1) but not CD45^-^ PD-L1^+^ (non-immune cells with PD-L1) cells compared with IgG controls (**Fig. 3p**). To assess if combination of PD-L1 with DTA-1 will be effective in tumors, we next examined this expression pattern in a published cohort of urothelial patients treated with anti-PD-L1 therapy^25^ (such a cohort is not available for PDAC). We applied genes highly expressed in DTA-1- treated compared to IgG control-treated 3275 tumors to stratify urothelial patients into DTA-1 sensitive-and resistant-like groups (see Methods section). DTA-1 sensitive-like patients survived longer (15.4 months) than DTA-1 resistant-like patients (7.9 months; **Fig. 4a**) with anti-PD-L1 treatment, as assessed using overall survival. Furthermore, the DTA-1 sensitive-like group was associated significantly (p=0.006) with complete and partial responses than stable and progressive diseases but not DTA-1 resistant-like group (**Fig. S8a(i))**. Interestingly, the DTA-1 sensitive-like group was significantly (p<0.0001) associated with >1% PD-L1 protein expression in immune cells by IHC (IC2 in **Fig. S8a(ii)**) and with inflamed immune phenotypes (**Fig. S8a(iii))**, mirroring 3275 DTA-1-treated tumors with increased PD-L1 expression in CD45^+^ immune cells compared with the DTA-1 resistant-like class. These results suggest that DTA-1 treatment responses may be improved when combined with anti-PD-L1 treatment, possibly due to release of PD-L1-associated T-cell exhaustion further boosting immune cell responses.

**Figure 4.**
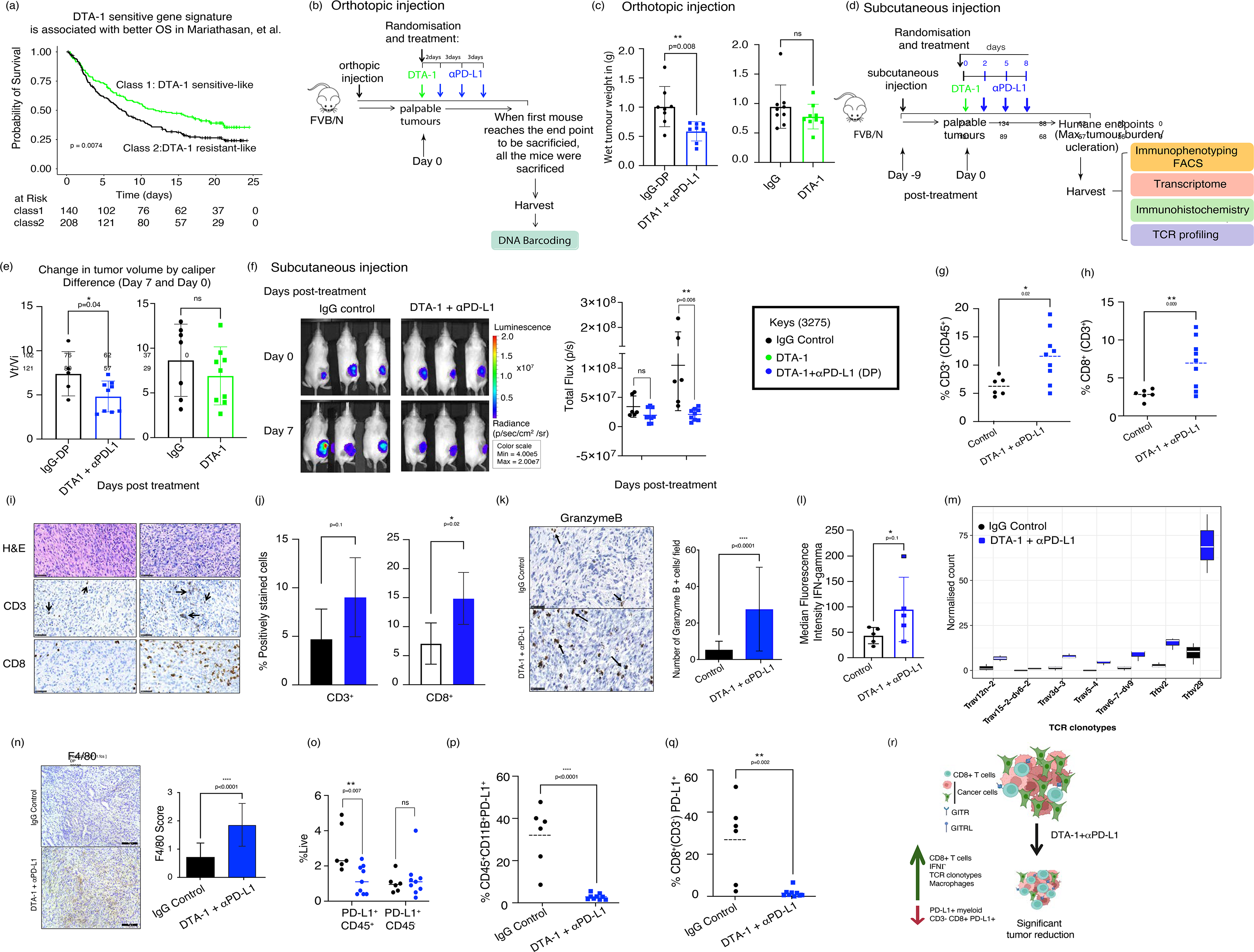
Improved response to DTA-1 with anti-PD-L1 combination treatment. (a) Kaplan-Meier curve (log-rank test for p value) showing prognostic difference between two groups, DTA-1 sensitive-like and resistant-like, of urothelial patients treated with anti-PD-L1 therapy. DTA-1-based groups were derived using gene signatures from DTA-1 and IgG treated 3275 mouse model (SAM analysis). (b) An orthotopic model-based schematic representing DTA-1+ anti-PD-L1 (αPD-L1) combination therapy and vehicle control in 3275 model. (c) A barplot showing the wet weight of orthotopic tumours in IgG (two different IgG clones corresponding to DTA-1 or combination of DTA-1 and αPD-L1), DTA-1 and DTA-1 + αPD-L1 treated groups. (d) A subcutaneous model-based schematic representing DTA-1 + αPD-L1 combination therapy and vehicle control in 3275 model. (e) A barplot showing the change in tumor volume between IgG control, DTA-1 and DTA-1 + αPDL1 treatments. Vi – tumour volume initially and Vt – tumour volume at day 7 after treatment. (f) Representative figure and quantitation of tumor (subcutaneous) volume as luminescence intensity in mouse tumours treated with IgG or DTA-1 + αPD-L1 combination. Luminescence was measured using IVIS® Lumina X5 system. (g-h) Flow cytometery data showing differential g) CD45^+^CD3^+^ and h) CD3^+^CD8^+^ populations in IgG vs. DTA-1 + αPD-L1 combination treatment. (i-j) IHC i) representative figure and j) quantitation of CD3^+^ and CD8^+^ T cells in 3275 mice treated with IgG or DTA-1 + αPD-L1. (k) IHC representative figure and quantitation of granzyme protein expression in 3275 mice treated with IgG or DTA-1 + αPD-L1. (l) ELISA results with quantitation of IFN-γ in 3275 mice treated with IgG or DTA-1 + αPD-L1. (m) Absolute count of TCRα/β clonotypes in IgG or DTA-1 + αPD-L1 combination treatment. (n) IHC showing F4/80 pan-macrophages in IgG or DTA-1 + αPD-L1 treated 3275 model. (o) Flow cytometry data showing differential CD45^+^ PD-L1^+^ and CD45^-^ PD-L1^+^ population in IgG vs. DTA-1 + αPD-L1 combination treatment. (p-q) Flow cytometry data showing differential p) CD45^+^CD11B^+^PD-L1^+^ and q) CD3^-^CD8^+^PD-L1^+^ populations in IgG vs. DTA-1 + αPD-L1 combination treatment. (r) Schematic summarizing microenvironment and cancer cell changes associated with DTA-1 + αPD-L1 combination treatment in 3275 mouse PDAC. Unpaired t test was done for p values for c, e-l,n-q.

To combat PD-Ll-mediated T cell exhaustion, we examined if combined DTA-1 and PD-Ll blockade (DP experiment) further reduced tumor burden and improved survival *in vivo* in the orthotopic and subcutaneous GITRL^high^GITR^high^ 3275 models. There was a significant reduction in tumor volume in DP treated compared with control mice in orthotopic 3275 model (**Fig. 4b-c** and **Fig. S7n**). Similarly, in subcutaneous model, there was a significant (p<0.05) reduction in tumor burden in the DP group (by caliper measurements) compared with IgG controls and a borderline significant reduction compared with DTA-1 monotherapy (**Fig. 4d-e**). Moreover, IVIS imaging showed significant reductions in subcutaneous tumor signal (**Fig. 4f**). The combination treatment significantly improved mouse survival compared to controls (**Fig. S8b**). This suggests that anti-PD-L1 therapy combination with DTA-1 is beneficial in 3275 mice.

Congruent with the change in tumor volume, the DP group showed dense CD3^+^ and CD8^+^ T cell infiltrates and modest, but border-line significant (p=0.1), upregulation of granzyme-B and IFN-γ protein levels (**Fig. 4g-l**; modest change in granzyme-B and IFN-γ protein levels because tumors were examined at an endpoint of mouse survival at which time the sensitive tumors have progressed). Consistently, there was a significant increase in seven TCRα/β clonotypes as assessed by gene expression analysis (**Fig. 4m**). These data suggest that combination DP treatment promotes favorable anti-cancer immune infiltrates.

Notably, there was a significant increase in pan-macrophages (positive for F4/80) compared with controls (**Fig. 4n**). In contrast, there was a significant reduction in PD-L1^+^ cells in the immune microenvironment (CD45^+^) but not the non-immune (CD45^-^) microenvironment showing overall reduction in total PD-L1 in the tumor (**Fig. 4o; Fig. S8c-d**). Specifically, DP treatment reduced PD-L1^+^ cells representing CD11B^+^ myeloid and CD3*^-^*CD8^+^ cells (potentially NK cells) compared with controls (**Fig. 4p-q**; but not the overall CD11B^+^ myeloid and CD3*^-^*CD8^+^ cells; **Fig. S8e-f**). However, there was no change in overall CD45^+^ immune and CD45^-^ non-immune cells reflecting no change in barcoded cancer cells, including cancer-initiating cells, as assessed by DNA barcoding technology (**Fig. S8g-k**). There was no change in FOXP3 expression (**Fig. S8l-m**). These results suggest that specific immune populations with PD-L1 expression were affected with the combination DTA-1+anti-PD-L1 treatment. **Fig. 4r** summarizes the events leading to an anti-tumor response mediated by combination DP therapy and the rationale for choosing anti-PD-L1 in combination.

#### Chemotherapy in combination with GITR agonist and anti-PDL1 further enhances efficacy

To further establish clinical relevance, we tested the combination of standard-of-care gemcitabine/nab-paclitaxel with DP immunotherapy in parental 3275 tumors (GADP) (**Fig. 5a**). There was a significant survival benefit with GADP compared with vehicle control (log rank p<0.001) (**Fig. 5b**), and an improved median survival benefit for GADP over DP (27 vs. 23 days; 14% improved survival). There was a reduction in tumor volume in GADP-and DP-treated tumors compared with control (significant; p<0.05; **Fig. 5c**).

**Figure 5.**
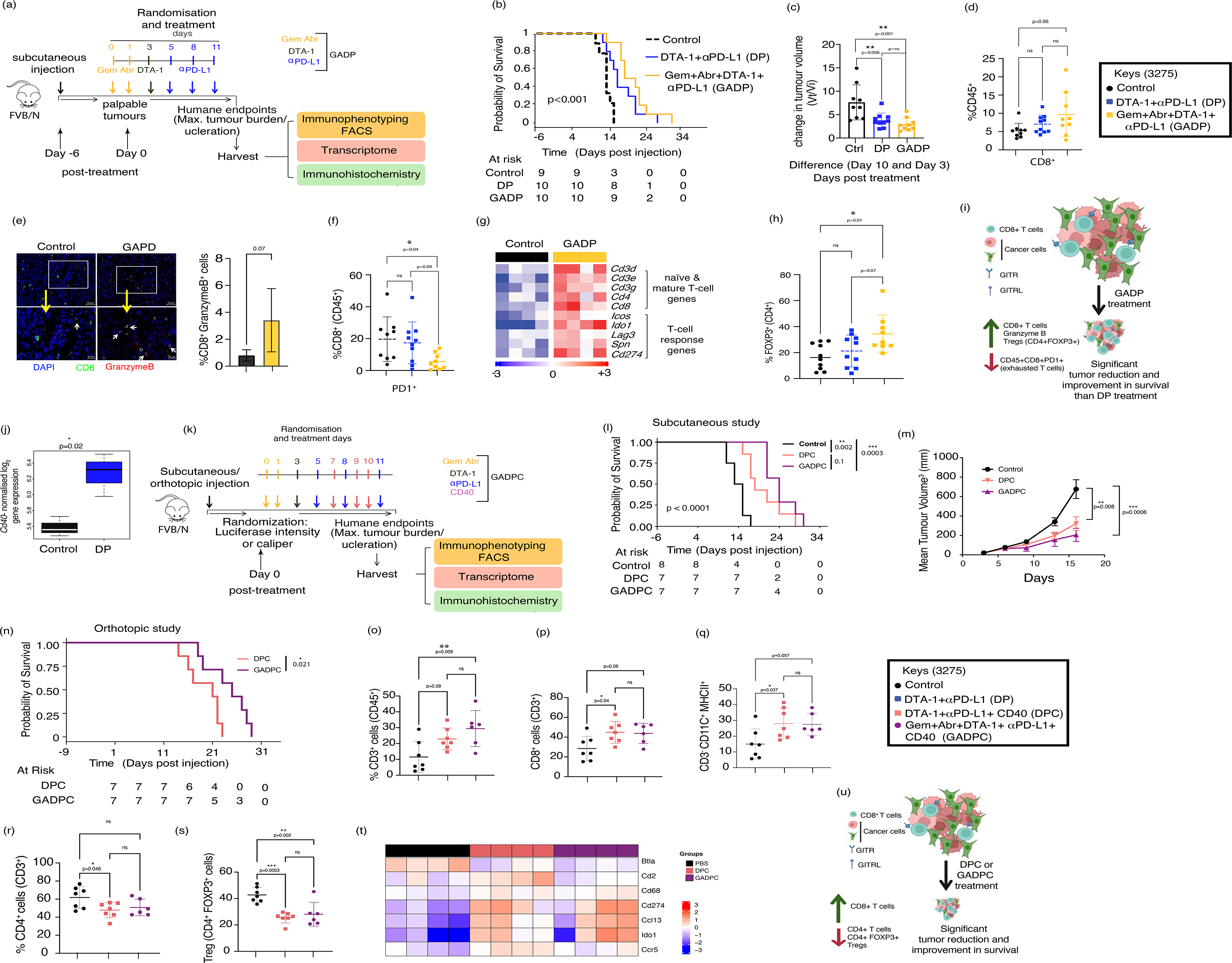
Combination of chemotherapy with GITR agonist, anti-PD-L1 and CD40 agonist therapies improved further response. (a) A schematic representing chemotherapy (gemcitabine and abraxane), DTA-1 and anti-PD-L1 combination therapy in 3275 model. (b) A Kaplan-Meier plot showing survival (log-rank test for p value) of mice treated with PBS, DTA-1 + anti-PD-L1 or chemotherapy + DTA-1 + anti-PD-L1 combination therapies. (c) A barplot showing change in tumor volume between PBS, DTA-1 + αPD-L1 or chemotherapy + DTA-1 + anti-PD-L1 combination therapies. (d) Plot showing change in CD45^+^CD8^+^ cells as measured by flow cytometry in PBS, DTA-1/anti-PD-L1 or chemotherapy/DTA-1/anti-PD-L1 combination therapies. e) Immunofluorescence and quantitation of CD8^+^ Granzyme^+^ T-cells in chemotherapy/ DTA-1/anti-PD-L1 combination therapies and PBS control. (f) Plot showing change in CD45^+^CD8^+^PD1^+^ exhaustive T-cells as measured by flow cytometry in PBS, DTA- 1/anti-PD-L1 or chemotherapy/DTA-1/anti-PD-L1 combination therapies. (g) Heatmap of immune genes differentially expressed between PBS and chemotherapy/DTA-1/anti-Pdl1 combination therapies. (h) Plot showing change in CD4^+^FOXP3^+^ exhaustive T regulatory cells as measured by flow cytometry in PBS, DTA-1/anti-PD-L1or chemotherapy/DTA-1/anti-PD-L1 combination therapies. (i) Schematic summarizing microenvironment and cancer cell changes associated with chemotherapy/DTA-1/anti-PD-L1 combination therapies in 3275 mouse PDAC. (j) Boxplots showing increased normalized log_2_ expression of *Cd40* gene in PBS and DTA-1/anti-PD-L1. (k) A schematic representing chemotherapy (gemcitabine and abraxane), DTA-1, anti-PD-L1 and CD40 agonist combination therapies in 3275 model. (l) A Kaplan-Meier plot showing survival (log-rank test for p value) of subcutaneous mice treated with PBS, DTA-1 + anti-PD-L1 + anti-CD40 or chemotherapy + DTA-1 + anti-PD-L1 + anti-CD40 combination therapies. (m) Plots showing mean tumor volume over days between PBS, DTA-1 + anti-PD-L1 + anti-CD40 or chemotherapy + DTA-1 + anti-PD-L1 + anti-CD40 combination therapies. (n) A Kaplan-Meier plot showing survival (log-rank test for p value) of orthotopic mice treated with DTA-1 + anti-PD-L1 + anti-CD40 or chemotherapy + DTA-1 + anti-PD-L1 + anti-CD40 combination therapies. (o-s) Plot showing change in (o) CD45^+^CD3^+^, (p) CD3^+^CD8^+^, (q) CD3^-^CD11C^+^MHCII^+^ APC, (r) CD3^+^CD4^+^, and (s) CD4^+^FOXP3^+^ (Treg) cells as measured by flow cytometry in PBS, DTA-1 + anti-PD-L1 + anti-CD40 or chemotherapy + DTA-1 + anti-PD-L1 + anti-CD40 combination therapies. (t) Differentially expressed genes representing key immune genes, specifically in T-cells in PBS, DTA-1 + anti-PD-L1 + anti-CD40 or chemotherapy + DTA-1 + anti-PD-L1 + anti-CD40 combination therapies. (u) Schematic summarizing microenvironment and cancer cell changes associated with DTA-1 + αPD-L1 + αCD40 or chemotherapy + DTA-1 + anti-PD-L1 + αCD40 combination therapies in 3275 mouse PDAC. Unpaired t test for p values for j and one-way Anova multiple comparison for c-f, h, m, o-s.

Investigation of immune responses elicited by GADP revealed a modest, but a border-line significant, increase in CD8^+^ T-cells in GADP tumors compared to control (**Fig. 5d**). Indeed, immunofluorescence analysis revealed higher expression of cytotoxic granzyme-B-expressing CD8^+^ T cells in the GADP group compared with vehicle control (**Fig. 5e**). CD8^+^PD-1^+^ cells were reduced (**Fig. 5f**), compared with DP treatment and controls, suggesting a reduction in exhausted T-cells in GADP tumors. Further, transcriptome analysis revealed that, compared with vehicle control, GADP treatment significantly upregulated T-cell genes (**Fig. 5g**): naïve and mature T cell genes (*Cd3d, Cd3e, Cdg, Cd4, Cd8*) and T-cell response (co-stimulatory and inhibitory) genes (*Icos, Ido1, Lag3, Spn*). In contrast, there was an increased CD4^+^FOXP3^+^ Tregs in GADP group compared to the DP treated and control samples (**Fig. 5h**). Overall, there was a significant improvement in response to GADP compared with vehicle control in 3275 PDACs but potentially resistance occurs via increased Tregs (**Fig. 5i**).

#### CD40 agonist to chemo-immunotherapy combination additionally improves therapy response

The addition of chemotherapy to DP was not significant in 3275 model (**Fig. 5c**). Hence, we sought to further improve therapy response when treated with chemotherapy by objectively selecting the next immunotherapy target in addition to DP treatment. Interestingly, another agonist CD40 showed increased expression in 3275 tumors treated with DP or GADP (**Fig. 5j and Fig. S9a**). Hence, we treated 3275 model with CD40 over DP (DPC) or GADP (GADPC) treatments (**Fig. 5k**). Remarkably, there was significantly (p<0.0001) improved survival in DPC and GADPC compared to control (**Fig. 5l**). This difference in survival reflected in tumor volume with a significantly (p<0.05) and more than 2.5-fold reduced mean tumor volume in DPC, that was even more pronounced in GADPC (**Fig. 5m and Fig. S9b**). Furthermore, we performed orthotopic injection of 3275 cells followed by GADPC, DPC or vehicle control. We observed significant difference in survival between DPC and GAPDC (also with vehicle; **Fig. 5n** and **Fig. S9c**). The modestly improved survival (although non-significant) with DPC relative to DP and GADPC relative to GADP was evident (**Fig. S9d**) when these results were combined with the previous results (**Fig. 5b**). This suggests that combination of chemotherapy with CD40 and DP improved the overall survival over DPC alone (**Fig. 5n and Fig. S9c**).

We further performed flow cytometry to analyze changes in immune cell population in DPC vs. GAPDC vs. control. The improved response to DPC and GAPDC were associated with increased CD3^+^ T-cells, CD8^+^ T-cells and CD3^-^CD11C^+^MHCII^+^ APC and reduced CD4^+^FOXP3^+^ Tregs (p<0.05; **Fig. 5o-q,s**). However, CD3^+^ CD4^+^ T cells were not changing between GADPC and control but significantly (p<0.05) reduced in DPC vs control (**Fig. 5r**). There was also enrichment of T-cell-specific genes in DPC and/or GADPC compared to control samples (**Fig. 5t**). This chemo-immunotherapy combination with anti-CD40 combination therapy further improved therapy response in 3275 model relative to the previous combinations with specifically reduced Tregs and increased CD8^+^ T cells (**Fig. 5u**).

#### Contrasting effect of DTA-1 in the GITRL^low^GITR^med^ PDAC model

The preceding data show that GITR-mediated T cell modulation is context dependent in terms of oligoclones within a heterogeneous tumor. Interestingly, while GITRL^high^GITR^high^ 3275 mice responded to DTA-1, the DTA-1-mediated immune effects were very different in the 7947 model (GITRL^low^GITR^med^), with weak GITRL expression and significantly worse survival (**Fig. 1m**). There was an increase in tumor burden on DTA-1 treatment compared with the IgG control (**Fig. 6a**). Furthermore, DTA-1 treatment significantly (p<0.05) downregulated CD45^+^, CD8^+^ (including GITR^+^ border-line significance), and FOXP3^+^ (both genes and proteins) and modest, but not significant, CD4^+^FOXP3^+^ T cells by flow cytometry and IHC (**Fig.6b-e; Fig. S10a**).

**Figure 6.**
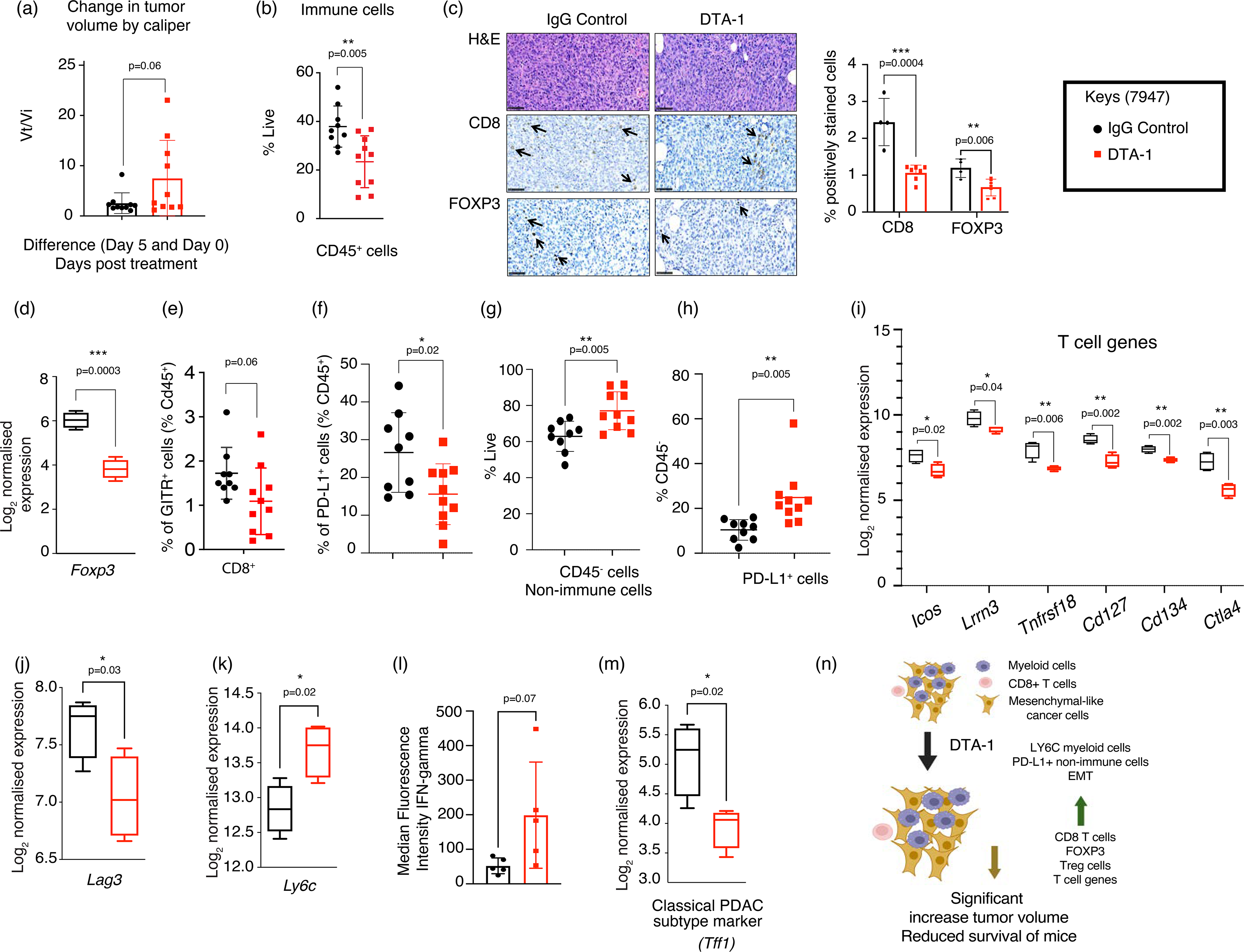
Context-specific resistance to anti-GITR in 7947 mouse model. (a) A barplot showing change in tumor volume between IgG and DTA-1treated 7947 model. (b) A dotplot based on flow cytometry showing CD45+ cells between IgG and DTA-1 treated 7947 model. (c) IHC (representative figure) and quantitation of CD8+ and FOXP3+ cells in 7947 mice treated with IgG or DTA-1. (d) A boxplot showing reduced Foxp3 gene expression in 7947 mice treated with IgG or DTA-1. (e-h) flow cytometry-based quantitation of e) CD45+ GITR+ CD8+ (cytotoxic T cells), f) CD45+ PD-Ll+, g) CD45- cells, h) CD45- PD-L1+ in 7947 mice treated with IgG or DTA-1. (i) A boxplot showing expression of T-cell and immune checkpoint genes in 7947 mice treated with IgG or DTA-1. (j-k) A boxplot showing j) Lag3 and k) Ly6c gene expression in 7947 mice treated with IgG or DTA-1. (l) A barplot showing IFN□ protein expression (Luminex) in 7947 mice treated with IgG or DTA-1. (m) A boxplot showing classical PDAC subtype marker gene, Tff1 in 7947 mice treated with IgG or DTA-1. (n) Schematic summarizing microenvironment and cancer cell changes associated with DTA-1 treatment in 7947 mouse PDAC with mesenchymal characteristics. Unpaired t test for p values for a-m.

In the 7947 model, while there was a significant reduction in immune microenvironment-specific PD-L1 expression (**Fig. 6f**) and no change in overall *Pdl1* gene expression (**Fig. S10b**), there was a significant increase in non-immune cells expressing PD-L1, consistent with increased tumor volume in contrast to 3275 model (**Fig. 6a, g-h; Fig. 3p**). Moreover, there was significant (p<0.05) downregulation of T-cell co-stimulatory markers (*Icos, Lrrn3, Tnfrsf18, Cd127, Cd134,* and *Ctla4*) and exhaustion markers (*Lag3*) (**Fig. 6i-j**). Of note, there was significant (p<0.05) upregulation of the myeloid-derived monocyte cell marker, *Ly6c* (**Fig. 6k**). Despite the reduction in cytotoxic T-cells and its co-stimulatory genes, there was a >2-fold increase, but border-line significant (p=0.07), in IFNγ protein expression upon DTA-1 treatment (**Fig. 6l**). The concomitant increase in *Ly6c* gene and IFNγ protein suggests an inflammatory response mediated by innate immune monocytes. Finally, there was significant downregulation of epithelial *Tff1* upon DTA-1 treatment (**Fig. 6m**), suggesting that immune infiltration may further modify the cancer cell phenotype (**Fig. 6n**). These results indicate that GITRL+GITR expression and DTA-1 therapy mediates context-dependent anti-tumor immune responses in different PDAC models with distinct immune populations that can be exploited for personalized therapy.

## Discussion

Immunotherapy is now considered the “fourth pillar” of cancer therapy. However, PDAC patients derive only marginal benefit from immune checkpoint blockade with anti-PD-1/PD-L1, CD40 and anti-CTLA4 agents.^1, 26^ An alternative strategy involves modulating cytolytic T cell activities in tumors via immunomodulatory therapies, specifically agonists.^27, 28^ There is increasing evidence from phase 1/2 studies that combined GITR agonism and checkpoint inhibition may be useful in patients with melanoma and other solid cancers. In a trial of the MEDI1873 GITR agonist, 42.5% of solid and neuroendocrine tumors had stable responses, including in 2 of 2 pancreatic neuroendocrine tumors.^29^ Given that ideally responses would be partial or complete, combination approaches, such as a bi-specific antibodies targeting GITRL and PD-1, are now being tested,^30^ but there remains the need to find other efficaciouscombination regimens for specific patient populations defined on the basis of robust biomarkers of response, and an improved understanding of the immune mechanisms underlying treatment outcome.

Here we stratified PDACs into four immune subgroups based on the expression of GITR and its ligand and validated our approach in mouse models receiving a GITR agonist in combination with PD-L1 blockade and standard of care chemotherapy. Zappasodi et al. reported that a reduction in Tregs or the ratio of effector T-cells to Tregs was a potential way to assess GITR agonistic efficacy in melanoma. However, to the best of our knowledge, there is no solution to stratifying patients for GITR agonists. Therefore, here we defined a GITRL/GITR score and tested it to stratify melanoma patients treated with adoptive T-cell therapy,^22^ a similar approach to boosting cytolytic T cells with GITR agonists.

We demonstrated the utility of the GITRL+GITR score in PDAC to demonstrate context-specific response of PDAC mouse models to GITR agonism (as mono and combination therapies). DTA-1 increased survival and decreased tumor burden in the GITRL^high^GITR^high^ 3275 model compared with models expressing lower levels of GITRL/GITR and was efficacious *in vitro*. Confirmation of the DTA-1-mediated increase in CD8^+^ T cells and PD-L1 expression *in vivo* and *in vitro* makes our models highly relevant for testing and validating immunotherapeutic agents. Systematic profiling of DTA-1-responsive 3275 tumors showed increased tumor infiltration with CD8^+^ effector T cells and an increase in granzyme-B and IFN-γ expression. Moreover, DTA-1 treatment was also associated with increased APP, immunoproteosome, and TCR changes. Despite the lack of FOXP3^+^ Treg depletion, there was a strong T-cell-mediated cytolytic response. This is consistent with other reports showing that changes in the Treg population are not the only indicator of GITR agonist responses.^13^

GITR agonist treatment may be associated with different cellular phenotypes. Although pathway enrichment analysis primarily showed EMT in controls but not after GITR agonist treatment, we observed a mixture of epithelial-and mesenchymal-like cells after DTA-1 treatment in the 3275 model. There was, therefore, considerable heterogeneity in this model, likely reflecting the clinical disease. By barcoding and identifying oligoclones in the 3275 model, we determined that the mesenchymal oligoclone is a persistent cell type primarily resistant to DTA-1, while epithelial clones showed acquired resistance to DTA-1 through PD-L1 expression. This epithelial oligoclone was more immunogenic *in vivo* than the mesenchymal clone, so other strategies will be required to target the mesenchymal clone effectively.

As demonstrated in other studies^15^ and results from our oligoclones, we observed T-cell exhaustion (increased *Pdl1* and *Lag3*) after one dose of DTA-1 in parental 3275 tumors *in vivo*. Multiple DTA-1 doses did not significantly improve cytolytic T-cell activity. Therefore, persistent T cell exhaustion mandates the use of combination therapy, and indeed combining DTA-1 with anti-PD-L1 therapy reinvigorated cytotoxic T-cell function and improved survival (**Fig. 3e-h; Fig. S8b**). This was associated with increased cytotoxic T and NK cells and M1 macrophage responses and decreased immunosuppressive PD-L1+ myeloid cells to induce a pro-inflammatory and anti-tumor microenvironment. Interestingly, dual targeting significantly reduced PD-L1-expressing CD45^+^ cells but not CD45^-^ cells, suggesting differential targeting of the immune and non-immune PD-L1-expressing compartments. This response to anti-PD-L1 treatment after DTA-1 in humans was further supported by a bioinformatics analysis stratifying urothelial cancer patients treated with anti-PD-L1 using DTA-1 response signatures from mice.

Even though the majority of PDACs are chemo-refractory, gemcitabine has remained standard-of-care. Our data, corroborated by other studies, suggest that combined chemo-immunotherapy is superior to immunotherapy alone. Protein expression data confirmed synergy between the two therapeutic modalities, as indicated by significantly increased CD8^+^ T cell infiltrates that were less exhausted, as indicated by reduced PD-1 expression compared with immunotherapy alone. These data, coupled with the observation of CD8 and granzyme-B co-immunostaining of tumor sections, indicate that the anti-tumor response increases due to the additional effect of chemo/immunotherapy within this model.

Importantly, context dependent GITR activity was evidenced by the distinctly different tumor behaviors of the GITRL/GITR low models. The 2334 model, with low expression of both GITRL and GITR, showed no response to anti-GITR treatment. By contrast, the GITRL^low^GITR^high^ (7947) model showed reduced survival and increased tumor burden in treated mice. The 7947’s phenotypes were associated with a significant reduction in T-cell-associated genes and decreased CD8^+^ T cell infiltration after DTA-1, supporting a pro-tumorigenic microenvironment. The data also highlight that, despite the inflammatory response in treated tumors, there was no tumor clearance; rather the non-immune cells (including cancer cells) increased, possibly through inflammation-induced tumor promotion. Hence, these results emphasize the importance of personalized selection of patients for immunotherapy.

In summary, we show that dual targeting with a GITR agonist and anti-PD-L1 therapy with or without chemotherapy of tumors expressing both GITRL and GITR is an effective therapeutic strategy, which can be further enhanced by rational selection and additional targeting of PD-L1 and CD40. Furthermore, using a rational approach, we pre-selected models displaying differential responses to DTA-1. This combined GITR agonism approach, also supported by the recent success of combination GITR immunotherapy in advanced chemo-refractory tumors, now needs testing clinically. This is the first report showing that GITRL/GITR expression can guide selection of patients for GITR agonist therapy, and that further combination with chemo-or other immunotherapies may enhance the therapeutic benefit for patients.

## Methods

### Cell lines

Three murine PDA cell lines from primary PDAC tumours (FVB/NCrl; Charles River, Alderley Park, UK) of female and male transgenic mice representing - *p48-Cre; LSL-Kras^G12D/+^; LSL-p53^R172H/+^* (KPC-p48) or *p48-Cre; LSL-Kras ^G12D/+^; Ink4a/Arf ^flox/flox^* (KIC) were obtained under material transfer agreement with Prof. Douglas Hanahan (Swiss Federale Institute of Lausanne; EPFL, Lausanne, Switzerland). The sub-lines or oligoclones from these lines were further developed using cloning cylinders (Sigma-Aldrich, Dorset, UK) as per the manufacturer’s protocol.

To generate luciferase expressing cancer cell lines, plasmid pCDH-CMV-MCS2-EF1-Hygro (System Biosciences Inc., Palo Alto, CA, USA) cloned with Luc2 open reading frame (ORF; Luc-plasmid) was obtained as a gift from Professor Sue Eccles (Institute of Cancer Research, London). Luc-plasmid was packaged with pPACKH1 plasmid mix using the protocol described by the manufacturer (System Biosciences Inc.). Cell lines were transduced with Luc-plasmid using Sigma^®^ MISSION^®^ ExpressMag^®^ Beads (Sigma-Aldrich) and selected under hygromycin antibiotic (Sigma-Aldrich). The cell lines were genotyped and the genetic background was confirmed (**Table S1**). All cell lines were mycoplasma screened before animal injections through a PCR-based method (Supplementary data).

### *in vivo* experiments

The mice with 5-6 weeks of age and female FVB/NCrl were purchased from Charles River Laboratories. All animal research was performed under the auspices of animal protocols approved by UK Home Office Regulations under the Animals Scientific Procedures Act 1986 and national guidelines (Project licence-P0A54750A). 0.5 x 10^6^ luciferase-transduced cells in 100 μl of Hank’s balanced salt solution (HBSS; Gibco, NY, USA) were injected subcutaneously in the right flank of female FVB/N immune competent mice (Charles River, UK). Animals were housed in specific pathogen-free rooms in autoclaved, aseptic micro-isolator cages with a maximum of five animals per cage. Tumour diameter was measured every 3 days with digital callipers and the volume was calculated using the formula, *Tumor* 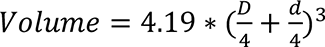 (D and d are large and small diameters, respectively). Tumour growth was also monitored weekly via luminescence emitted by the luciferase transfected cells upon injection of d-luciferin substrate, using the IVIS® Lumina. X5 (PerkinElmer, Beconsfield, UK). Randomisation into respective treatment groups was done when the tumours reached an average tumour volume of 50 mm^3^. *Treatment regime:* single dose 10 or 20 mg/kg DTA-1 (Catalog number; Cat; #BE0063; BioXCell) or rat IgG2b isotype control (Cat - #BE0090, BioXCell); *for combination treatment:* single dose 20 mg/kg DTA-1 or isotype control followed by 3 doses of 10 mg/kg anti-PDL1 (obtained in collaboration Bristol Meyers Squibb; B7-H1) or isotype control (mouse IgG2a, Bristol Meyers Squibb) with a 2-day interval, 2 days post DTA-1 administration; *for chemo-immuno combination treatment:* single dose 120 mg/kg Gemcitabine (Royal Marsden Hospital) or PBS control, followed by single dose 120 mg/kg Abraxane (Royal Marsden Hospital) or human albumin control (Sigma) the next day. The same regime of DTA-1 and anti-Pdl1 was followed as described earlier after 1 day and 3 days post-Abraxane injection respectively. All compounds were administered intra-peritoneally. For survival analysis, mice were sacrificed when they reached maximum tumour burden (average of D from tumor volume calculation and d> 15 mm or d> 17 mm) or an ulceration of grade 4. Tumors were harvested for further experiments.

For orthotopic injection of cancer cells into pancreas, cancer cells were resuspended in 1:1 Hank’s balanced salt solution (HBSS; Gibco, NY, USA) and Matrigel (Corning, Weisbaden, Germany), after a wash with PBS,. For tumor cell injection, mice underwent gaseous anaesthesia and their surgical site was shaved and disinfected. The pancreas was exposed for orthotopic injection through a small incision under aseptic conditions. A total of 50,000 luciferase-transduced cancer cells were injected in 20uL final volume into the pancreas. The incision was sutured. Tumour growth was monitored by palpating three times per week. Studies were terminated when the first animal in the study got sick or reached tumour burden. Upon the first animal becoming unwell or tumor reaching maximum volume when palpated, the study was terminated.

### Flow cytometry analysis of mouse tumours

Tumours transported in MACS tissue storage solution were enzymatically and mechanically dissociated into single cell suspensions using MACS mouse tumour dissociation kit following manufacturer’s instructions and filtered through 70 μm cell strainers. Cell viability was tested using Trypan blue dye (Invitrogen). 2 x 10^6^ viable cells were stained and incubated at 4°C for 20 minutes for cell surface markers listed in **Table S2.** Fixable dye was used in the panel to negatively select viable cells during analysis. Intracellular staining was performed after fixation and permeabilisation of the cells using FOXP3 Fixation/Permeabilisation Solution kit (BD Biosciences, Frankin Lakes, NJ, USA) using manufacturer’s instruction. Analysis was performed on BD FACSAria^TM^ II (BD Biosciences). Flow cytometry data was analysed using BD FACS Diva software (BD Biosciences) and FlowJo portal (FLOWJO, Ashland, OR, USA).

### Generation of cell line-derived three-dimensional (3D) organotypic cultures

3275 mouse cancer cells, derived from GEM tumours, were used to generate 3D organotypic cultures (adopted from our previously published protocol, Vlachogiannis et al^31^). Cells were seeded in 120 μl of growth factor reduced Matrigel (Corning, Flintshire, UK) in 24-well plates. Matrigel was left to solidify by incubating at 37°C for 20 minutes and was then overlaid with 500 μl of mouse organoid growth media (advanced DMEM supplemented with 2mM L-glutamine, 0.01% BSA, 1x B27 additive, 1x N2 additive, 50 ng/ml EGF, 100 ng/ml Noggin and 500 ng/ml R-Spondin1). Cells were incubated at 37°C with 5% CO_2_ and media was refreshed every 2 days.

### Isolation, activation, and co-culture of mouse splenocytes for in vitro immunotherapy

Spleens from non-tumour bearing FVB/NCrl mice (Charles River, Alderley Park, UK) were harvested and transported in advanced DMEM supplemented with 10% FBS and 2% Penicillin-Streptomycin (PS; Gibco, NY, USA). Mechanical disruption of spleens was performed using sterile scalpels and collected in 50 ml falcons. After two washes in phosphate buffered saline (PBS; Gibco, NY, USA), the contents were digested in 2x TrypLE (Gibco, NY, USA) in PBS containing 1mM ethylenediamine-tetraacetic acid (PBS-EDTA) solution containing 20 μl DNase for 1 hour at 37°C. Undigested tissue particles were filtered through a 70 μm cell strainer and the single cell suspension was subjected to Red Blood Cell (RBC) lysis using ACK (Ammonium, Chloride, Potassium) lysis buffer (Gibco, NY, USA) for 3-5 minutes on ice. Cells were washed in cold PBS, re-suspended in media and counted. Splenocytes were activated by seeding at a density of 2×10^6^ cells in T cell media (advanced DMEM supplemented with 10% FBS, 2mM L-glutamine (Gibco, NY, USA), 1% PS, 2mM Concanavalin A (Sigma, St. Louis, USA) and 50 IU recombinant IL-2 (Peprotech, London, UK) in 24-well plates and incubated at 37°C with 5% CO_2_ for 48 hours. 1 x 10^6^ activated (Concanavalin A + IL2) splenocytes were incubated in conditioned media from cancer cell organotypic cultures and treated with 20 μg/ml rat anti-mouse DTA-1 (BioXcel Therapeutics, New Haven, CT, US) or rat isotype IgG2b (BioXcel Therapeutics). Cells were collected after 72 hours, and viable cells were stained with fluorophore-conjugated antibodies for 20 min at 4°C. Cells were subjected flow cytometry analysis.

### Ribonucleic acid extraction from frozen mouse tumours

Flash frozen tumours were homogenised in Precellys beaded tubes (Bertin Technologies^TM^, Montigny-le-Bretonneux, France) containing 700 μl of TRIzol^TM^ using a homogeniser. Total RNA from the tissue digest was extracted using miRNeasy FFPE Kit (Qiagen^TM^, Manchester, UK) as per manufacturer’s instructions. The yield was quantified using NanoDrop^TM^ (Thermo Fisher Scientific^TM^, Delaware, USA).

### NanoString nCounter Transcriptomic Profile – low-cost custom-designed mouse panel

100 ng of total mRNA was used to run a custom designed mouse panel (72 genes, **Table S2**) on the nCounter® Max analysis system (NanoString Technolgies, CA, USA) as per the manufacturer’s instructions. Raw data was processed, normalised and log2 transformed using nSolver 3.0 software.

### Targeted RNA sequencing

100ng RNA was subjected probe-based TempO-seq mouse whole transcriptome expression profiling assay (Bioclavis, Clydebank, Scotland). Sequencing and data analysis were done according to the service providers’ protocols. Bioclavis provided a table of raw count data, which was normalised using DESeq2 and was used for further downstream analysis.

### Immunoblotting

Flash frozen mouse tumours were homogenised in Precellys beaded tubes (Bertin Technologies^TM^, Montigny-le-Bretonneux, France) containing 100 μl of cold NP40 lysis buffer (50mM Tris-HCL, 150mM sodium chloride, 1% NP40) and phosphatase and protease inhibitors (Pierce, Life Technologies Corporation, CA, USA) using a tissue homogeniser (Bertin Technologies^TM^, Montigny-le-Bretonneux, France). Protein concentrations were determined using Bicinconinic acid kit (BCA; Pierce, Life Technologies Corporation, CA, USA) and 20 μg of protein was fractionated onto 12% sodium dodecyl sulphate-polyacrylamide gels. Blotting was performed on polyvinylidene difluoride (PVDF) membranes for 1 hour at 100 V. Membranes were blocked in PBS containing 5% BSA + 0.12% Tween-20 for 1 hour followed by probing with primary antibodies, overnight at 4°C. The membranes were washed and incubated with anti-mouse/rabbit secondary antibodies conjugated to horseradish peroxidase the following day. Immunoreactive bands were visualised on Licor Odyssey Fc (Cambridge, UK) using enhanced chemiluminescence (ECL, Merck Millipore, Darmstadt, Germany). β-actin was used a loading control.

### Immunohistochemistry

Tumours were fixed in 10% neutral buffered formalin for 48 hours and embedded in paraffin. Consequently, 4 μm thick serial sections were cut, deparaffinised in xylene, processed in graded alcohol, and rehydrated in water. One section was stained with haematoxylin and eosin for histological analysis. For single and multiplex immunostaining, antigen retrieval was performed using a pressure cooker method in the appropriate antigen retrieval solution (Citrate buffer, pH 6.0 or EDTA buffer, pH 9.0) for 10 minutes. Slides were allowed to cool in the solution and blocked with 3% Bovine Serum Albumin and Normal Goat Serum, where appropriate, for 1 hour at RT. Slides were then incubated with primary antibody at optimised antibody dilution overnight at 4°C. The slides were then washed thrice in wash buffer (Tris buffered saline with 0.025% Tween) and blocking of endogenous peroxidase was done using peroxidase block (Dako) for 15 minutes. For single immunostaining, after performing three washes, the slides were incubated in appropriate secondary antibody for 1 hour at RT. Protein expression was visualised using DAB (Dako Real Envision kit) for 3-5 minutes. Sections were counterstained with hematoxylin and mounted after dehydration in graded alcohol and xylene. For multiplex immunostaining, sections were first stained with anti-mouse CD8 primary antibody and incubated at 4°C overnight followed by goat anti-rat Alexa Fluor^TM^ 555 conjugated secondary antibody for 1 hour. Antibody stripping was performed before adding subsequent primary antibody by microwaving the sections for 5 minutes at maximum power. This was followed by staining with Granzyme B primary antibody and goat anti-rabbit Alexa Fluor 633 conjugated secondary antibody. The nuclei were stained with DAPI and the slides were mounted with VectaShield mounting media (Vector Labs). Appropriate positive and negative controls were used in all runs. Primary and secondary antibodies used in the study are listed in **Table S2**. Whole stained sections were scanned using Nanozoomer-XR C12000 for chromogenic staining and Zeiss Axio Scan Z1 for fluorescent staining. Representative images were taken using Zeiss LSM700 confocal microscope. Antigen expression was scored using Definiens image analysis software (Tissue studio version 4.4.3).

### Mouse Granzyme B ELISA

Flash frozen mouse tumours were lysed in Precellys beaded tubes (Bertin Technologies) containing 100 μl of NP40 lysis buffer containing a protease and phosphatase inhibitor cocktail (Invitrogen). The lysates were centrifuged at 2000 rpm for 10 min at 4°C. Supernatants were transferred to fresh pre-cooled 1.5 ml tubes (Eppendorf). Protein was diluted 1:15 and estimation was done using BCA kit (Pierce, Life Technologies Corporation, CA, USA) as per manufacturer’s instructions. 50 μg protein per sample was used to perform mouse Granzyme B Platinum ELISA (Thermo Fisher Scientific #BMS6029) and the protocol was followed as per the product manual.

### Mouse IFN-gamma Singleplex cytokine assay

20 μg protein per sample was used for the IFN-gamma mouse Procartaplex^TM^ Simplex High Sensitivity kit (Thermo Fisher Scientific) and the protocol was followed as per the product manual. IFN-gamma protein expression was analysed using Luminex xMAP technology on Luminex^TM^ MagPix200 (Thermo Fisher Scientific).

### Cellular DNA barcoding and NGS sequencing

The cellular DNA barcoding, next generation sequencing (NGS), and data analysis were performed as described in Aalam et al^32^. Briefly, cells were transduced with lentiviral EGFP_index 5 DNA barcode library at predetermined dilutions (< 37% efficiency). Two days later, viable (DAPI-) barcoded (EGFP+) single cells were purified by FACS (BD FACSAria^TM^ II; BD Biosciences) and injected in mice. Genomic DNA was extracted from 10 mg homogenized fresh frozen tumors using DNeasy® Blood & tissue kit (Qiagen). Subsequently, NGS sequencing libraries were prepared from 1 µg of genomic DNA spiked with known ‘spike-in’ controls by introducing Illumina adaptors and 5-bp-long index sequences using Q5® High-Fidelity 2X Master Mix (NEB). The barcode amplification was verified in parallel polymerase chain reaction (PCR) reactions. The NGS sequencing libraries were multiplexed, spiked with 10% commercially prepared PhiX library (Illumina) and sequenced as 100 base single end reads on an Illumina HiSeq 2500 using TruSeq Rapid SBS sequencing kit version 1 and HCS version 2.0.12.0 data collection software (Illumina, San Diego, CA). Base-calling is performed using Illumina’s RTA version 1.17.21.3. (Illumina) as described elsewhere.

The CIC calculator described elsewhere^32^ was subsequently implemented to extract barcode sequences from the raw FASTQ files, filter noise, determine fractional read value (FRV) thresholds based on known spike-in controls, and absolute clone counts for subsequent analyses. The samples for the barcoding was selected based on tumor burden.

### Datasets processed and analyzed

Publicly available PDAC datasets used were from ICGC RNAseq and microarray, The Cancer Genome Atlas (TCGA), GSE71729 and GSE62452. In addition, pan-cancer TCGA dataset was also used. Immunotherapy treated publicly available datasets, include GSE100797 and Mariathasan et al.^25^ Except for the immunotherapy datasets, normalized data from the publications were used.

### Differential gene expression analysis

Statistical Analysis of Microarrays (SAM^33^; using siggenes R package^34^) or t test was used for differential gene expression analysis.

### DTA-1 response genes and patient immunotherapy dataset analysis

SAM-based differential gene expression analysis was performed between DTA-1 treated and control 3275 tumors to identify two gene sets upregulated (high expression) in DTA-1 treated and control tumors. The DTA-1-high gene set was used to derive DTA-1-sensitive-like and DTA-1-resistant-like classes of melanoma patients from Mariathasan et al.^25^ (immunotherapy dataset). This was performed by calculating mean scores for each sample based on DTA-1 treated genes, followed by determining the optimal cut-point (threshold) using overall survival and mean scores for patient samples. A similar classification of control genes into high and low classes of patients was performed.

### Statistical analyses

GENE-E (from GenePattern^35^) was used for heatmap representations. Paired or unpaired t test, Kruskal-Wallis, Chi-square or Multiple comparison ordinary one-way Anova test was used to compare groups of samples. All other statistical analyses, including survival, were performed using Graphpad Prism 8 (for Windows, GraphPad Software; La Jolla, CA, USA, www.graphpad.com), XLSTAT 2019 or R packages.

## Contribution Statement

A.S., C.R., K.D., and N.S. conceived the idea. A.S., C.R., K.D., and P.V.L. designed the experiments. C.R., K.D., P.V.L. Y.I., M.M.A., H.Ps., and A.S. performed the experiments. D.C. supported the study. N.K. provided DNA barcoding reagents, helped design the barcoding experiments, and analyzed its associated data. A.S., K.D., and C.R. wrote the manuscript. A.S., and N.S. supervised the study. All the authors critically reviewed the manuscript.

## Acknowledgements

We would like to acknowledge BMS for both partial funding of this work and for providing anti-PD-L1 antibody for use in these experiments. We thank Prof. Douglas Hanahan, Dr. Stephan Wullschlegar and Dr. Mélanie Tichet, Swiss Federale Institute of Lausanne; EPFL, Lausanne, Switzerland for kindly providing mouse cell lines and tumours. We thank Prof. Alan Melcher from the Centre for Translational Immunotherapy, ICR for critically reviewing the manuscript and Mr. Adrian Larykerd for helping with TCR analysis. We thank Dr. Florence Raynaud, Ms. Melanie Valenti and Cancer Therapeutics Unit, ICR for kindly supplying KPC mouse colonies. We thank Dr. Ian Titley and Mrs. Louise Howell (Institute of Cancer Research) for supporting with flow cytometry and microscopy-based imaging analysis. We thank Dr. Elisa Fontana, Ms. Melanie Herpels, Mr. Aswanth Harish Mahalingham and other members of the Sadanandam Lab for their support.

## Supplementary Figures

**Supplementary Figure 1. Response to anti-PD1 therapy in PDAC mouse and expression of immunomodulatory genes in patient PDACs.** (a) Kaplan-Meier curve showing survival (log-rank test) of 3275 PDAC mouse model treated with anti-PD1 or IgG control. (b-d) Heatmaps showing clustering of patient PDAC samples (ICGC cohort; n=70) with immunomodulatory genes, b) ICOS:ICOSL, c) OX40 (TNFSF4):OX40L (TNFRSF4), d) TIGIT:PVR, and T-cell and immune checkpoint genes. (e) IHC showing expression of GITRL protein in epithelial and immune cell compartments of mouse 3275 model.

**Supplementary Figure 2. Single cell analysis of *GITR* and *GITRL* in human and mouse cancer samples.** (a) Uniform Manifold Approximation and Project (UMAP) plot showing single cell distribution of *GITR* and *GITRL* expression in different cell types from treatment naïve PDAC samples. (b) Dot plot showing expression of *GITR*, *GITRL*, T-cell receptor and immune checkpoint inhibitor genes in single cells from breast cancer. Data source: Single Cell Portal from the Broad Institute. (c-d) Single cell *Gitr* and *Gitrl* expression in (c) dot and (d) t-distributed stochastic neighbor embedding (tsne) plots from *Pdx-Cre; LSL-Kras^G12D/+^; LSL-p53^R172H/+^* (KPC) mouse model.

**Supplementary Figure 3. *GITRL*/*GITR* expression in patient PDAC, pan-cancer and melanoma immunotherapy-treated samples.** (a-g) Boxplots showing expression of a) *GITR*, b) *GITRL*, c) *CD4*, d) *FOXP3*, e) *CD8A*, f) *PDL1*, and g) *LAG3* in four sub-groups of *GITR*/*GITRL* expression from ICGC PDAC (n=70) dataset. Kruskal-Wallis test was used for p values. (h) A heatmap showing significant (p<0.2 by Wilcoxon Test) enrichment of various immune profiles in *GITRL*^high^*GITR*^high^ vs. *GITRL*^low^*GITR*^low^ groups in TCGA patient PDAC samples. (i-j) Heatmaps showing the genes from a-g) in four other i) PDAC (n=545) and j) pancancer (n=9,077) datasets. (k-m) Assessment of (k) *GITRL*/*GITR* score based on gene expression data, (l) overall survival (log-rank test was used for p value), and (m) tumor response from melanoma patient samples (n=25; GSE100797) treated with adotive T-cell therapy between *GITRL*^high^*GITR*^high^ and *GITRL*^low^*GITR*^low^ groups. (n) A Kaplan-Meier survival curve showing prognosis and response to GITR agonist (20 mg/Kg) and its PBS control in PDAC GEM model. (o) A representative figure showing scoring of GITRL expression in mouse tumours from IHC.

**Supplementary Figure 4. Gating strategy for FACS analysis.**

**Supplementary Figure 5. Response to GITR agonist therapy *in vitro* and *in vivo* in mouse models.** (a) A schematic showing an *in vitro* indirect coculture system, where activated splenocytes were treated with conditioned media from mouse 3275 organoids (cancer cells grown in three-dimension using Matrigel). The activated splenocytes were treated with DTA- 1 or IgG, followed by flow cytometry analysis for immune cell types and markers. (b) 3275 organoids in 5x and 10x magnifications. (c) Abundance of CD8+ T cells by flow cytometry from 3275 co-culture system treated with DTA-1 or control IgG. (d-f) A Kaplan-Meier survival curves (log-rank test for p value) showing prognoses and response to DTA-1 therapy *in vivo* in syngeneic subcutaneous models (3275, 2334, 7947) of PDAC treated with GITR agonist (at different concentrations) and its control IgG. (g) A Kaplan-Meier survival curve (log-rank test for p value) showing prognosis and response to therapy *in vivo* in syngeneic orthotopic model (0787) of PDAC treated with GITR agonist (20mg/kg) and its controls - IgG and PBS.

**Supplementary Figure 6. GITR agonist response and its associated cellular and molecular changes in 3275 PDAC mouse model.** (a) A schematic representation of *in vivo* orthotopic 3275 PDAC model based GITR agonist response (interventional) evaluation, along with barcoding analysis. A bar plot showing wet tumor weight of orthotopic 3275 PDAC model treated with GITR agonist or IgG. (c-e) Various plots from subcutaneous 3275 PDAC model treated with GITR agonist or IgG showing c) *Cd8a* gene expression, d) CD45^+^ CD4^+^ cells from flow cytometry, and e) *Tff1* gene expression in 3275 mouse model treated with GITR agonist or IgG control. (f-h) Enrichment analysis showing increased antigen presentation and processing in tumors treated with GITR agonist and IgG control – f) Gene set enrichment analysis (GSEA^36^), and g-h) Pathview (a Bioconductor package^37^). (i) Boxplots showing expression of T-cell genes in 3275 tumors treated with GITR agonist and IgG control. (j) A Kaplan-Meier survival curve (log-rank test for p value) showing prognosis and response to therapy *in vivo* in syngeneic subcutaneous 3275 PDAC model treated with multiple doses of GITR agonist and its control IgG. (k) Barplot showing change in tumor volume in syngeneic models of PDAC treated with multiple doses of GITR agonist (at different concentrations) and its control IgG. Unpaired t test was performed for p values from b-e, i and k. Log-rank test was performed for j.

**Supplementary Figure 7. Heterogeneity and response to GITR agonist in 3275 mouse model**. (a) H&E showing heterogeneous regions in 3275 subcutaneous tumours. (b) A plot showing CD45^-^ cells in GITR agonist or IgG control treated 3275 subcutaneous model. (c) Electrophoretic gel of amplified PCR of barcoded DNA samples of treated and control mouse tumors. (d) Heterogeneity in 3275 cell line *in vitro* showing culturing of different oligoclones with distinct morphology. Magnification is 100μm. (e) Representative IHC figures showing how scoring of E-cadherin was done. Magnification is 100μm. (f-g) Bar plots showing IHC quantification of g) E-cadherin and h) N-cadherin. (h) A schematic showing an *in vitro* direct coculture system, where splenocytes and (two-dimensional) cancer cells were co-cultured. Magnification is 100μm. The co-culture systems were treated with DTA-1/DTA-1+αPD-L1 (DP) or IgG, followed by separating adherent (cancer) and non-adherent (splenocytes) and immunoblotting for different protein expression. (i) Immunoblot showing expression of cleaved caspase-3 in parent, and oligoclone #1, #4 and #5 treated with DTA-1 or IgG control. β-actin was used as a loading control. (j-k) Quantitation of PD-L1 expression from live cells from subcutaneous tumors after treated with DTA-1 or IgG using flow cytometry. (l) Quantitation of PD-L1 expression from *in vitro* co-culture from Supplementary Figure 5a-b using flow cytometry. (m) Immunoblot showing expression of cleaved caspase-3 and PD-L1 in parent, and oligoclone #1, #4 and #5 treated with DTA-1 + αPD-L1 treatment or IgG control using flow cytometry. (n) A barplot showing the wet weight of orthotopic tumours in DTA-1 and DTA-1 + αPD-L1 treated groups in 3275 model. Unpaired t test was done for p values for b, f, g, k, l and m.

**Supplementary Figure 8. Response to anti-PD-L1 in combination with GITR agonist in tumor.** (a) Bar plots showing association of class 1: DTA-1 sensitive-like and class 2: DTA-1 resistant-like urothelial carcinoma with (i) immunotherapy response based on Response Evaluation Criteria in Solid Tumors (RECIST) v1.1-criteria (CR – complete response; PR – partial response; SD – stable disease; and PD – progessive disease), (ii) PDL1 expression on immune cells (IC) (scored as IC3 representing ≥10% PD-L1 expression in ICs; IC2 - ≥5% but <10% PD-L1; IC1 - ≥1% but <5% PD-L1; and IC0 - <1% PD-L1), and (iii) immune subgroups – desert, excluded and inflamed. IC scores, RECIST and immune subgroups were represented as defined Mariathasan et al.^25^ DTA-1-based groups were derived using gene signatures from DTA-1 and IgG treated 3275 mouse model (see Methods). (b) Kaplan-Meier survival plot (log-rank test for p value) showing survival of mice from 3275 model treated with d) DTA-1+αPD-L1, or IgG or PBS controls. (c) Violin plot showing differential M2 macrophages in tumors treated with DTA-1 + αPD-L1 or IgG control (represented as mean expression scores). (d-e) FACS data showing quantitation of Pdl1 expression in tumors treated with DTA-1 + αPD-L1or IgG control. (f-g) Quantitation of g) CD3-CD8+ and h) CD11b+ live cells in tumors treated with DTA-1 + αPD-L1 or IgG control. (h-i) Quantitation of h) CD45+ and i) Cd45- live cells in tumors treated with DTA-1 + αPD-L1 or IgG control. (j-l) DNA barcoded results representing estimated total barcoded j) cells/tumor of tissue, k) clone size per tumor, and l) total clone-initiating cell (CIC) frequency in orthotopic tumors treated with DTA-1 + αPD-L1 or IgG control. P value refers to paired t test. (m-n) Plots showing quantitation of FOXP3 m) cells by FACS and n) log_2_ normalized gene expression in tumors treated with DTA-1 + αPD-L1 or IgG control. Unless represented, all the other p values were generated using paired t-test.

**Supplementary Figure 9. Response to DTA-1, anti-PD-L1 and CD40 agonist in combination with chemotherapy in 3275 tumor.** (a) A boxplot showing Cd40 gene expression changes between control and GADPC treated 3275 tumors. (b) A barplot showing mean tumor volume changes between DPC, GADPC and control treated mice. (c) Kaplan-Meier survival plot (log-rank test for p value) showing survival of orthotopic mice from 3275 model treated with DPC, GADPC and control treated mice. (c) Kaplan-Meier survival plot (log-rank test for p value) showing survival of mice from 3275 model treated with DP, DPC, GADP, GADPC and controls treated mice (combined results from two independent experiments). Unpaired t test for p value for a and one-way Anova multiple comparison for b.

**Supplementary Figure 10. Immune changes DTA-1 treated 7947 model.** (a-b) Change in a) T regulatory and b) *Pdl1* (*Cd274*) gene expression in subcutaneous 7947 model treated with DTA-1 and control.

## Supplementary Tables

**Supplementary Table 1. Genotyping results for the mouse cell lines used.** Genotyping was done by the company, Transnetyx

**Supplementary Table 2. Antibodies used for immunohistochemistry and flow cytometry.**

**Supplementary Table 3. List of genes used in mouse NanoString Analysis**

