## Supplementary Figures for "Context-specific GITR agonism potentiates anti-PD-L1 and CD40-based immuno-chemotherapy combination in heterogeneous pancreatic tumors"

Supplementary Figure 1

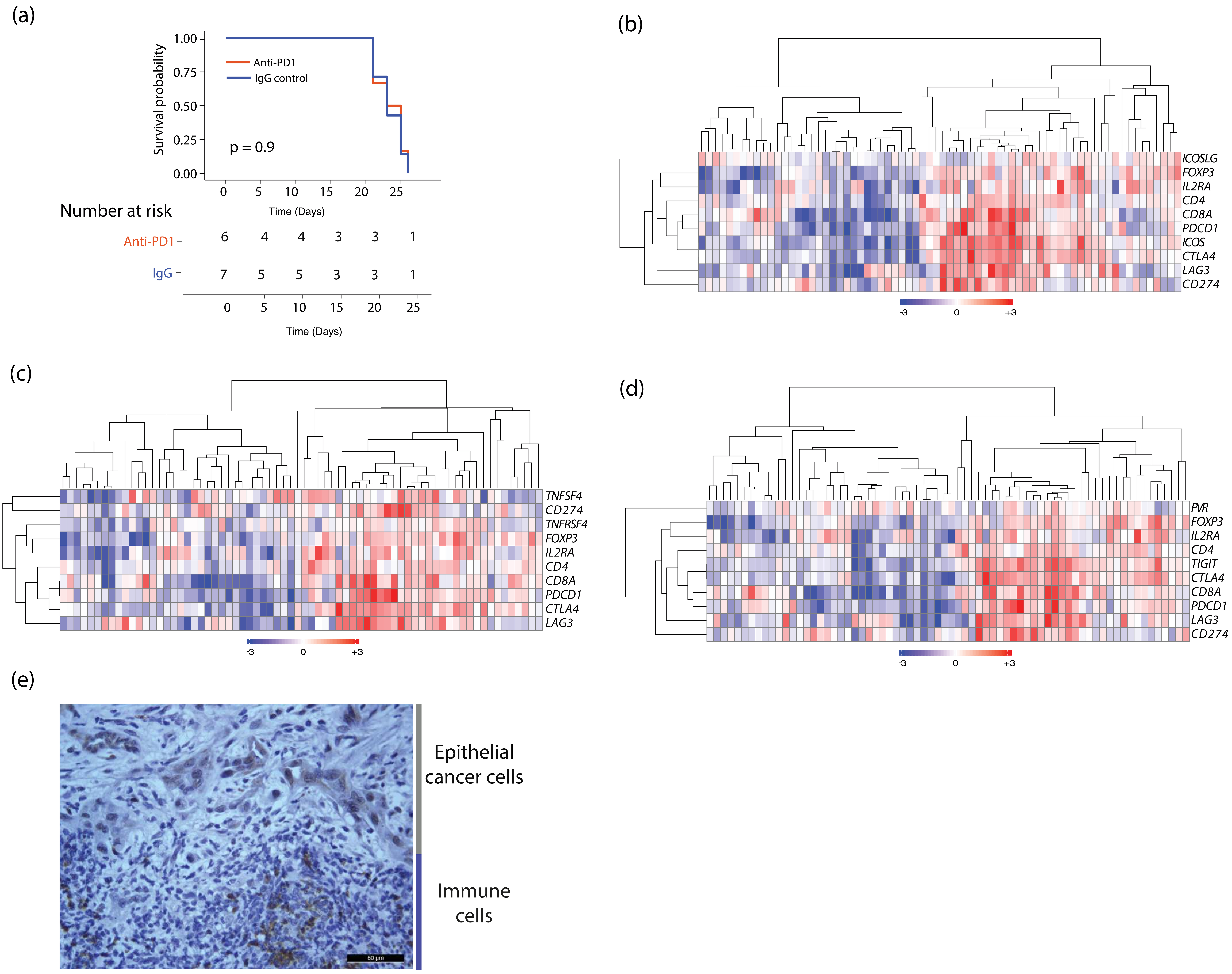

Supplementary Figure 2

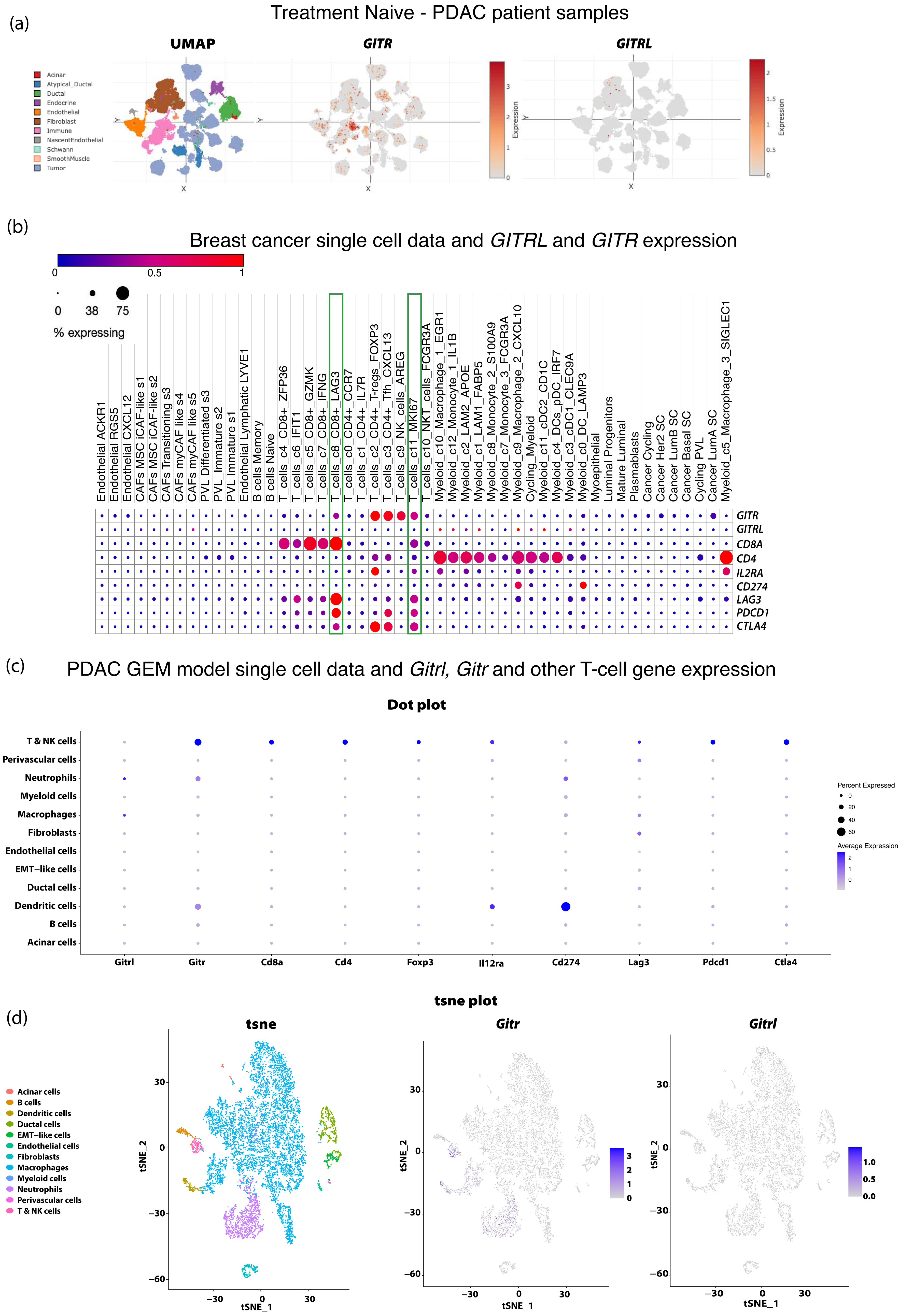

**Supplementary Figure 3**

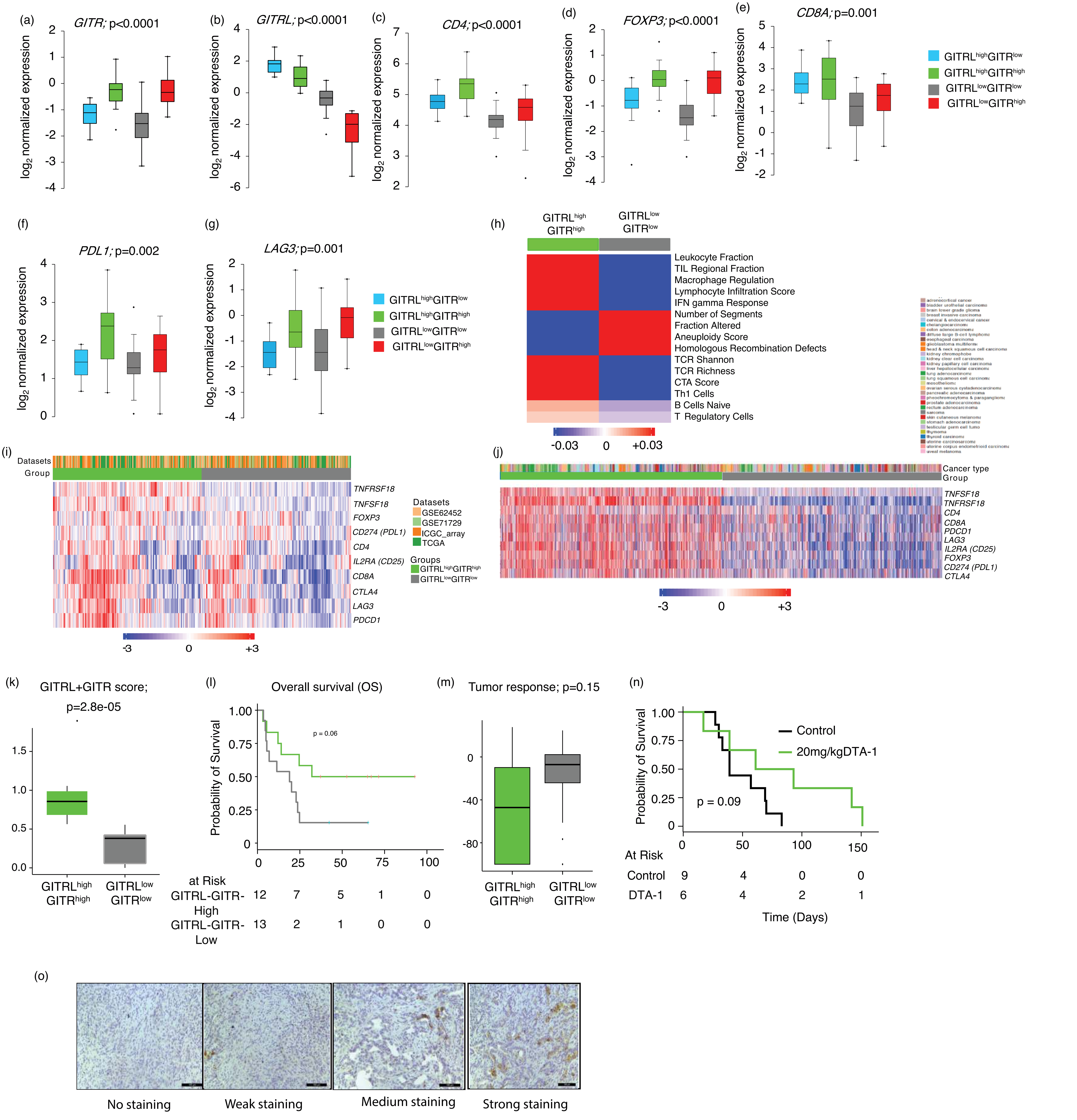

### Supplementary Figure 4

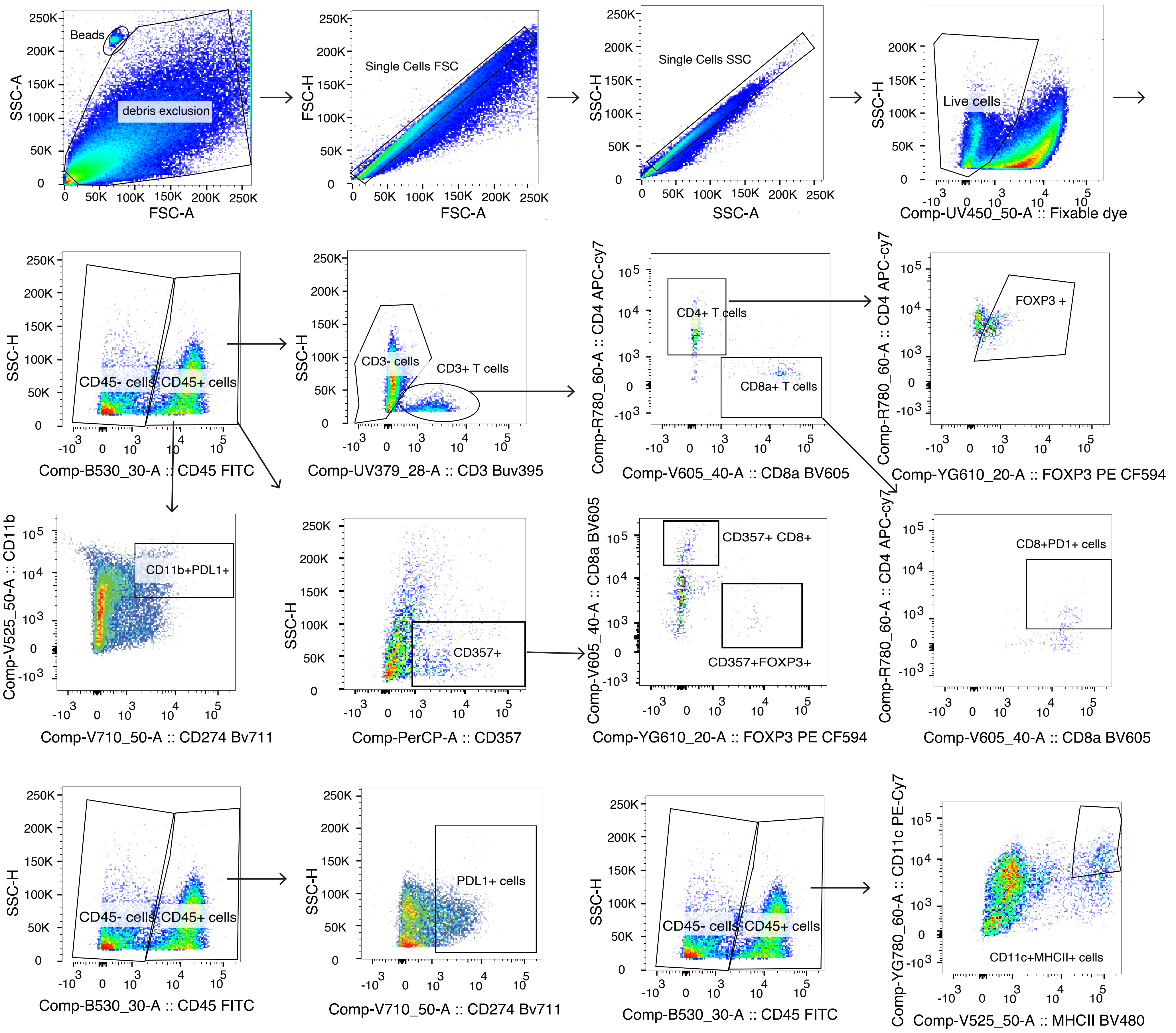

**Supplementary Figure 5**

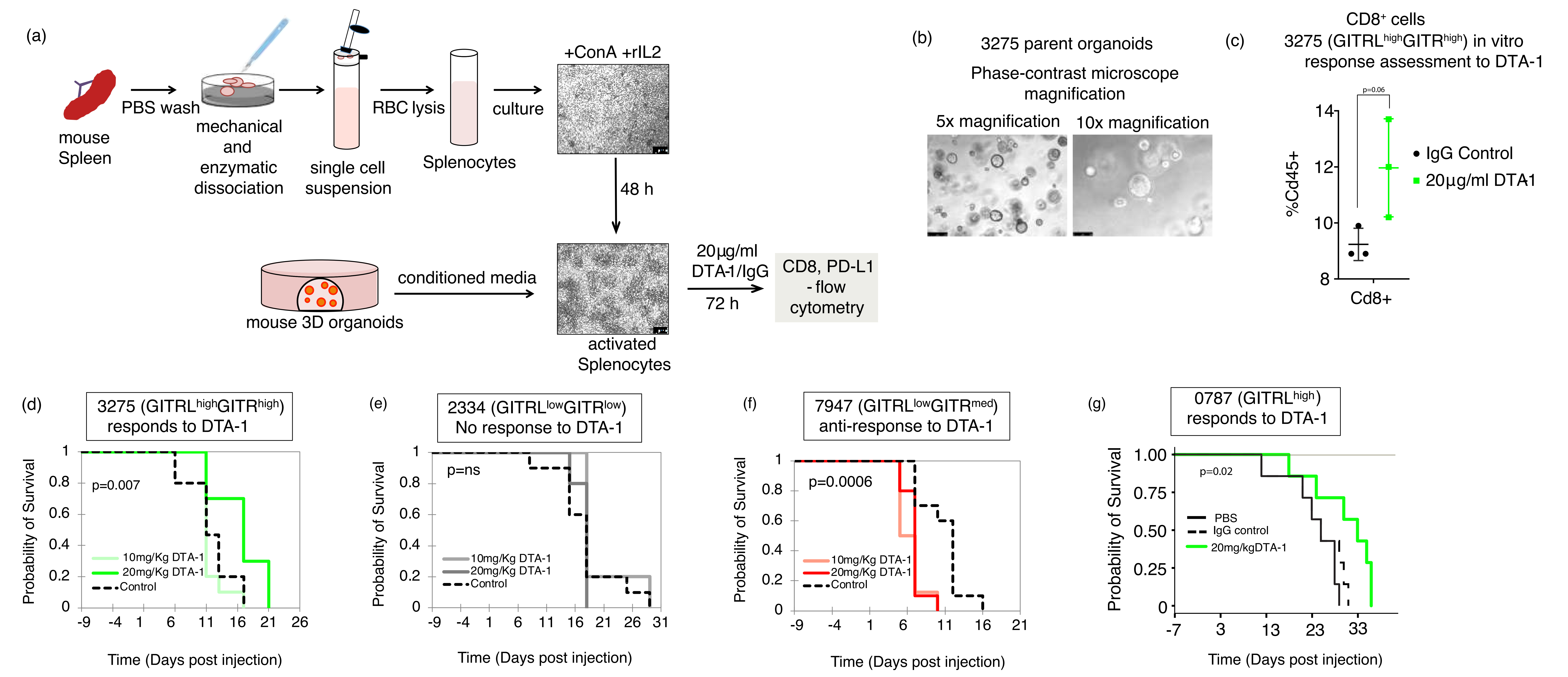

**Supplementary Figure 6**

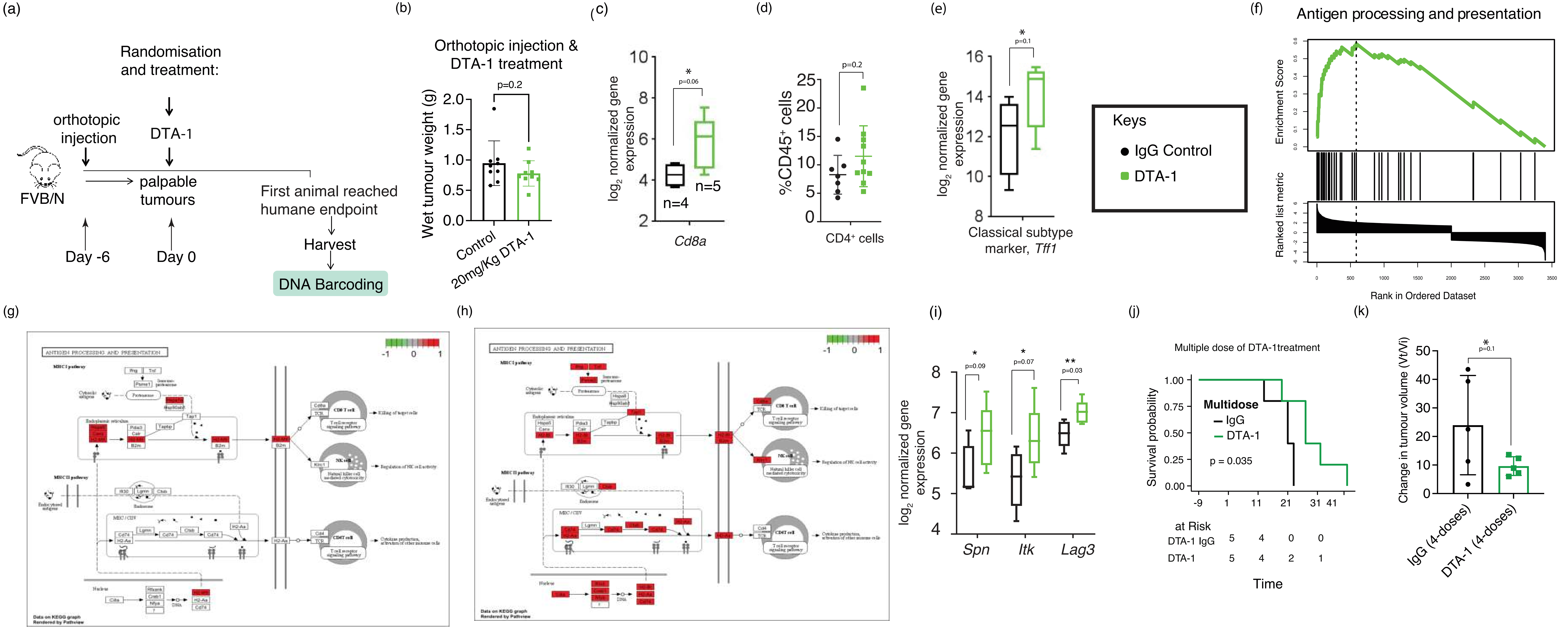

Supplementary Figure 7

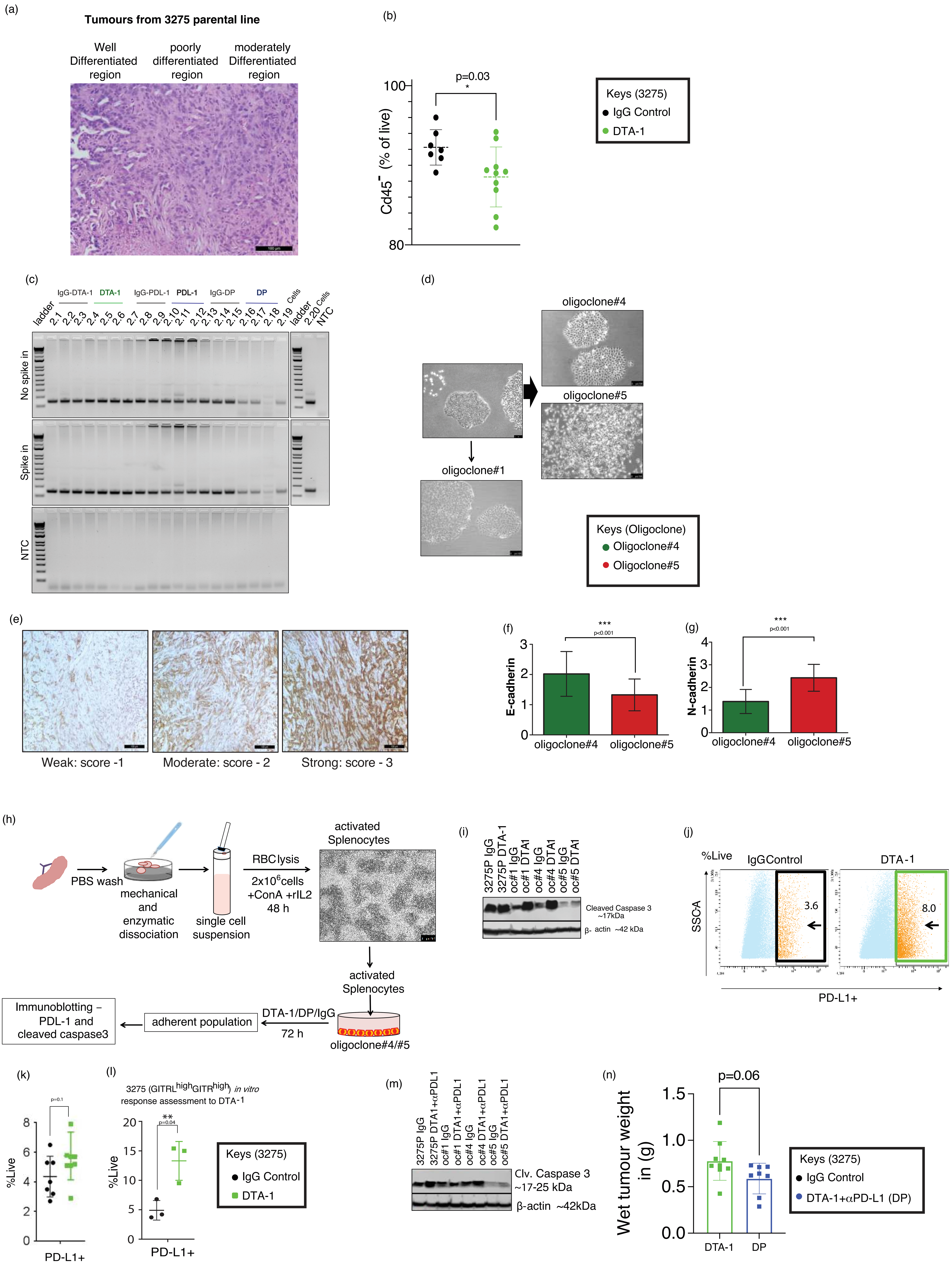

#### Supplementary Figure 8

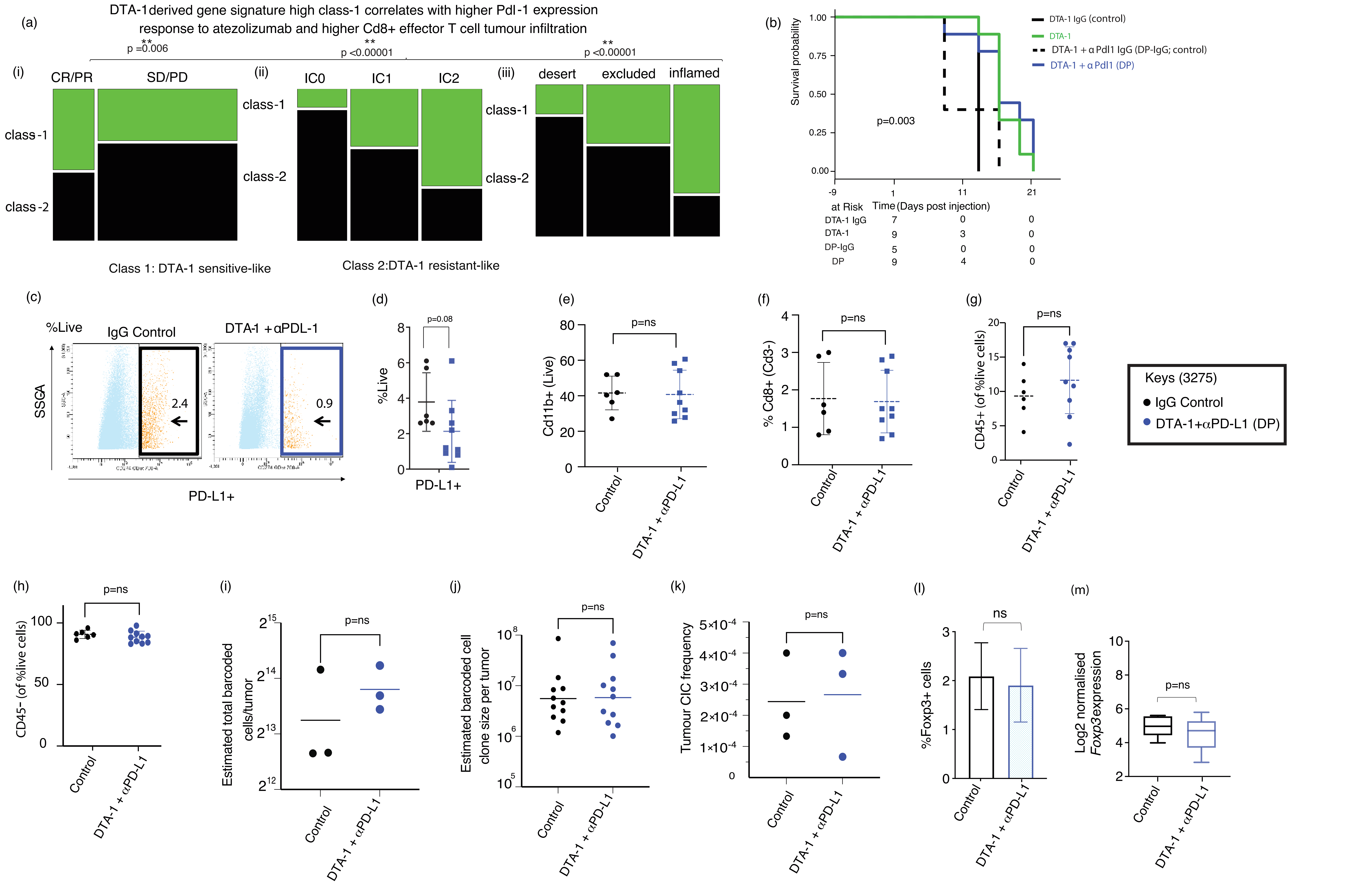

**Supplementary Figure 9**

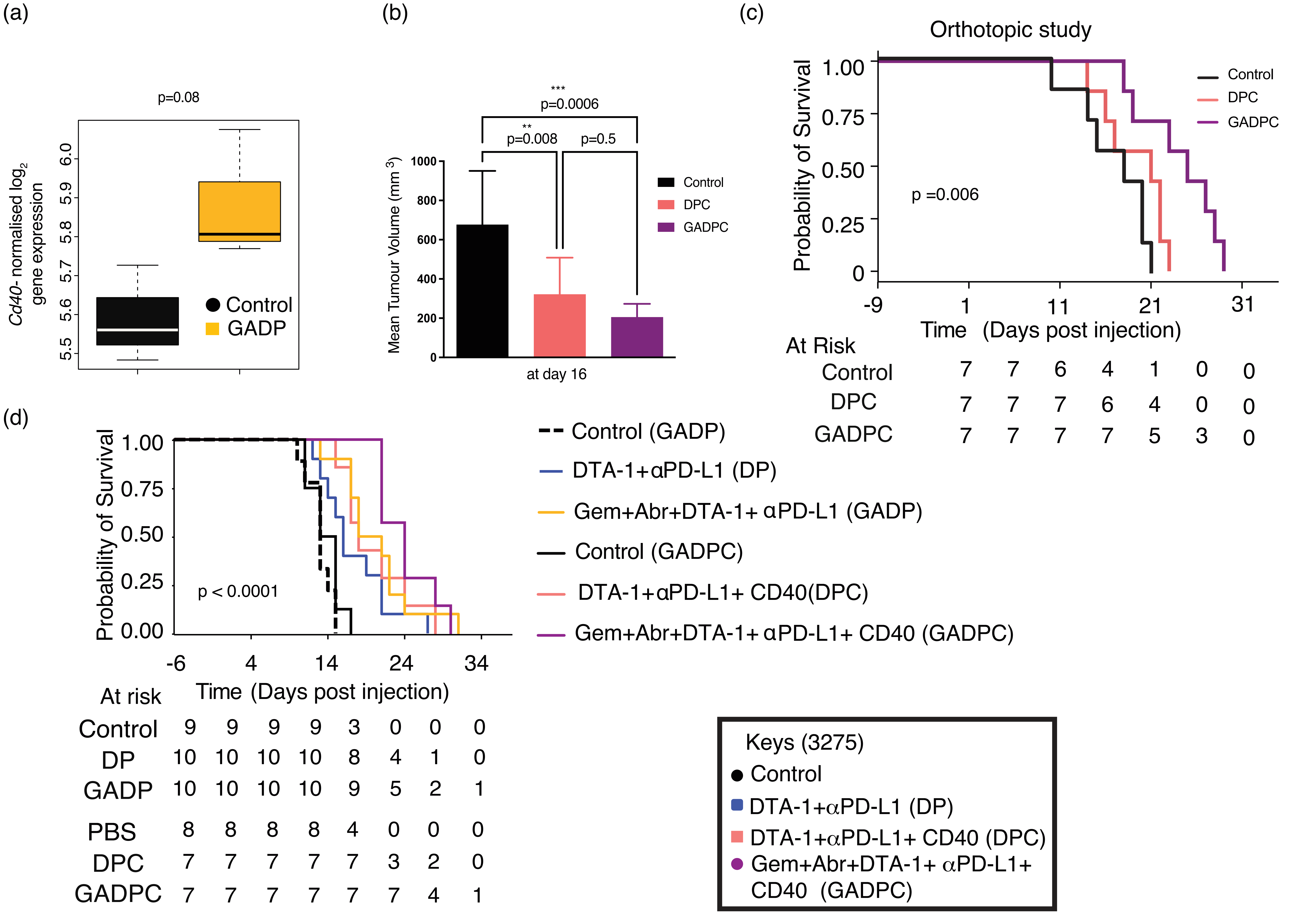

**Supplementary Figure 10**

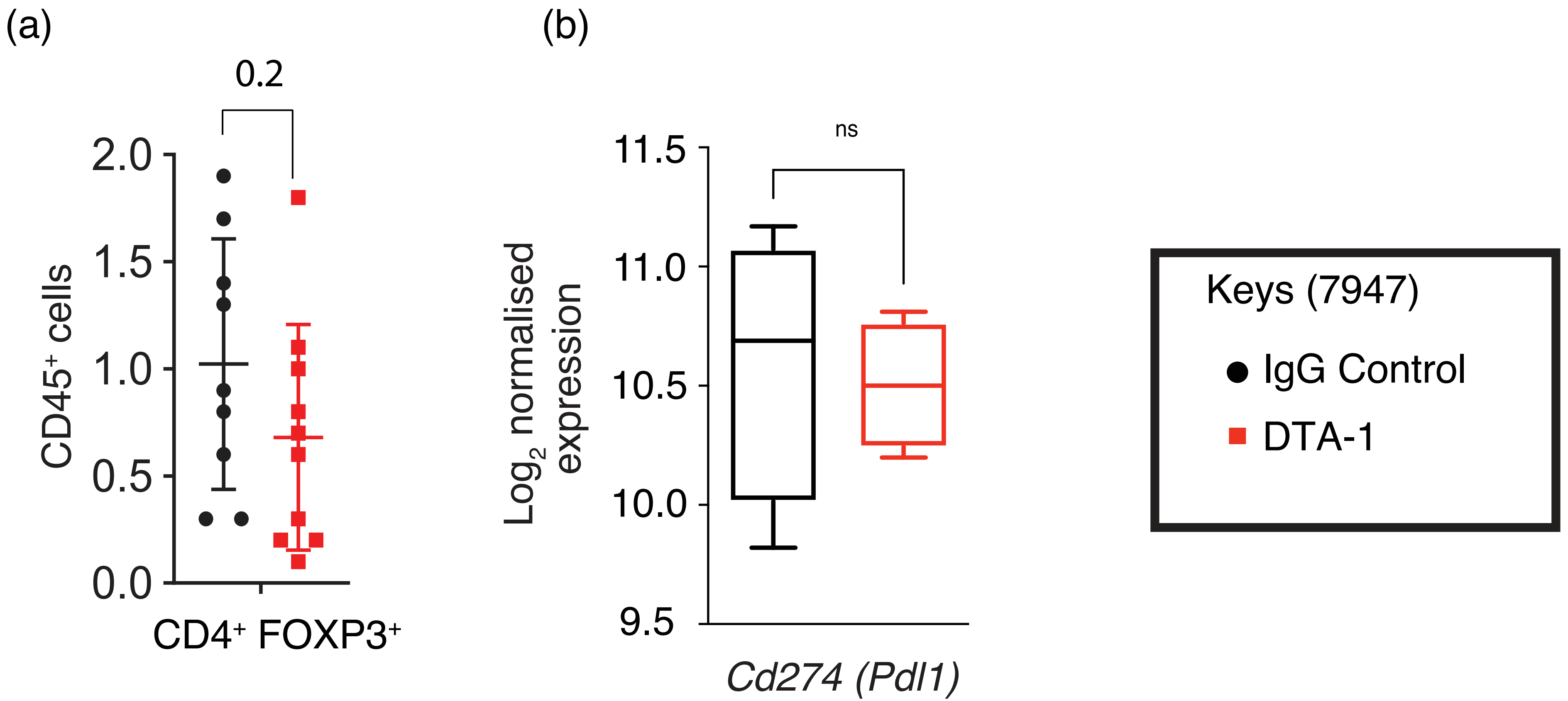
