## Supplementary Tables for "Context-specific GITR agonism potentiates anti-PD-L1 and CD40-based immuno-chemotherapy combination in heterogeneous pancreatic tumors"

**Supplementary Table 1. Genotyping results for the mouse cell lines used. Genotyping was done by the company, Transnetyx**

| **Samples** | **CRE** | **Kras G12D mutation** | **Trp53-1 mutation** | **Cdkn2a-2 Flox** | **Cdkn2a-2 wild-type** | Mouse model |
|---|---|---|---|---|---|---|
| 2334 with Luciferase | + | +- | ++ | - | + | KPC-P48cre |
| 3275 with Luciferase | + | +- | ++ | - | + | KPC-P48cre |
| 7947with Luciferase | + | +- | -- | - | - | KIC-P48cre |
| Oligoclone #1 | + | +- | ++ | - | + | KPC-P48cre |
| Oligoclone #4 | + | +- | ++ | - | + | KPC-P48cre |
| Oligoclone #5 | + | +- | ++ | - | + | KPC-P48cre |

KPC - p48-Cre; LSL-KrasG12D/+; LSL-p53R172H/+ model

KIC - p48-Cre; LSL-Kras G12D/+; Ink4a/Arf flox/flox

**Supplementary Table 2. Antibodies used for immunohistochemistry and flow cytometry**

| REAGENTS or RESOURCES | SOURCE | IDENTIFIER |
|---|---|---|
| **Antibodies for Immunohistochemistry** | | |
| CD3-epsilon (D4V8L) Rabbit Anti-Mouse mAb 1:100 | Cell Signalling Technologies | #99940S |
| CD4 (4SM95) Rat Anti-Mouse mAb 1:250 | eBioscience | #14-9766-82 |
| CD8a (4SM15) Rat Anti-Mouse mAb 1:250 | eBioscience | #14-080882 |
| CD31 (SZ31) Rat Anti-Mouse mAb 1:75 | Dianova | DIA310 |
| FOXP3 (D6O8R) Rabbit Anti-Mouse mAb 1:100 | Cell Signalling Technologies | #12653 |
| Granzyme B (D6E9W) Rabbit Anti-Mouse mAb 1:100 | Cell Signalling Technologies | #46890S |
| GITRL Rabbit Anti-Mouse pAb 1:100 | Abcam | ab203387 |
| Goat Anti-Rabbit Secondary | Dako | P0448 |
| Goat Anti-Rat Secondary | Vector Labs | PI-9401 |
| Goat Anti-Rabbit Alexa Fluor 488 | Invitrogen | A-11008 |
| Goat Anti-Rat Alexa Fluor 555 | Invitrogen | A-21434 |
| Goat Anti-Rabbit Alexa Fluor 633 | Invitrogen | A-21070 |
| Goat Anti-Rabbit Tyramide Signal Amplification Kit 488 | Invitrogen | B40943 |

**Antibodies for Flow Analysis**

| REAGENTS or RESOURCES | SOURCE | IDENTIFIER |
|---|---|---|
| CD45 (30-F11) FITC Anti-Mouse mAb 1:200 | BD Biosciences | #561088 |
| CD3 (17A2) Alexa Fluro 532 Anti-Mouse mAb 1:100 | Invitrogen | #58-0032-82 |
| CD4 (GK1.5) APC-H7 Anti-Mouse mAb 1:200 | BD Biosciences | #560264 |
| Cd8a (53-6.7) BUV805 Anti-Mouse mAb 1:200 | BD Biosciences | #564920 |
| CD274 (MIH5) BV711 Anti-Mouse mAb 1:200 | BD Biosciences | #563369 |
| CD279 (J43) BV421 Anti-Mouse mAb 1:200 | BD Biosciences | #562584 |
| CD357 (DTA-1) PE-Cy 7 Anti-Mouse mAb 1:200 | BD Biosciences | #558140 |
| CD11b (M1/70) BV480 Anti-Mouse mAb 1:200 | BD Biosciences | #566149 |
| FOXP3 (MF23) PE-CF594 Anti-Mouse mAb 1:200 | BD Biosciences | #562466 |
| Granzyme (NGZB) PerCP-eFluor 710 Anti-Mouse mAb 1:200 | Invitrogen | #46-8898-80 |

**Supplementary Table 3 - List of genes used in mouse NanoString Analysis.**

| Gene Name | Accession # | Class Name |
|---|---|---|
| Atp10b | NM_176999. | Endogenous |
| Btla | NM_0010377 | Endogenous |
| Cav1 | NM_007616. | Endogenous |
| Ccl13 | NM_011333. | Endogenous |
| Ccr5 | NM_009917. | Endogenous |
| Cd127 | NM_008372. | Endogenous |
| Cd134 | NM_011659. | Endogenous |
| Cd2 | NM_013486. | Endogenous |
| Cd25 | NM_008367. | Endogenous |
| Cd252 | NM_009452. | Endogenous |
| Cd274 | NM_021893. | Endogenous |
| Cd31 | NM_008816. | Endogenous |
| Cd37 | NM_007645. | Endogenous |
| Cd3d | NM_013487. | Endogenous |
| Cd3e | NM_007648. | Endogenous |
| Cd3g | NM_009850. | Endogenous |
| Cd4 | NM_013488. | Endogenous |
| Cd4/80 | NM_009855. | Endogenous |
| Cd79a | NM_007655. | Endogenous |
| Cd79b | NM_008339. | Endogenous |
| Cd8 | NM_0010811 | Endogenous |
| Cftr | NM_021050. | Endogenous |
| Cks2 | NM_025415. | Endogenous |
| Clec5a | NM_0010386 | Endogenous |
| Cma1 | NM_010780. | Endogenous |
| Colec12 | NM_130449. | Endogenous |
| Csf3r | NM_0012526 | Endogenous |
| Ctla4 | NM_009843. | Endogenous |
| Cxcl13 | NM_018866. | Endogenous |
| Elf3 | NM_0011631 | Endogenous |
| Erbb3 | NM_010153. | Endogenous |
| Fcer1a | NM_010184. | Endogenous |
| Foxp3 | NM_054039. | Endogenous |
| Foxq1 | NM_008239. | Endogenous |
| Gzmk | NM_008196. | Endogenous |
| Hmmr | NM_013552. | Endogenous |
| Icos | NM_017480. | Endogenous |
| Ido1 | NM_008324. | Endogenous |
| Itk | NM_010583. | Endogenous |
| Klrc2 | NM_010653. | Endogenous |
| Klrg1 | NM_016970. | Endogenous |

| | | |
|---|---|---|
| Krt14 | NM_016958. | Endogenous |
| Lag3 | NM_008479. | Endogenous |
| Lgals4 | NM_010706. | Endogenous |
| Lrrn3 | NM_010733. | Endogenous |
| Ly-6G | NM_0013104 | Endogenous |
| Ly-6c | NM_010741. | Endogenous |
| Ly9 | NM_008534. | Endogenous |
| Pdcd1 | NM_008798. | Endogenous |
| Pnliprp2 | NM_011128. | Endogenous |
| Ptf1a | NM_018809. | Endogenous |
| Sele | NM_011345. | Endogenous |
| Slamf1 | NM_013730. | Endogenous |
| Slc16a1 | NM_009196. | Endogenous |
| Spn | NM_0010378 | Endogenous |
| Tal1 | NM_011527. | Endogenous |
| Tff1 | NM_009362. | Endogenous |
| Tff3 | NM_011575. | Endogenous |
| Tmem45b | NM_144936. | Endogenous |
| Tnfrsf18 | NM_009400. | Endogenous |
| Tnfsf18 | NM_183391. | Endogenous |
| blk | NM_007549. | Endogenous |
| cd68 | NM_009853. | Endogenous |
| Eef1g | NM_026007. | Housekeeping |
| Hdac3 | NM_010411. | Housekeeping |
| Nubp1 | NM_011955. | Housekeeping |
| Oaz1 | NM_008753. | Housekeeping |
| Polr2a | NM_009089. | Housekeeping |
| Sap130 | NM_172965. | Housekeeping |
| Sdha | NM_023281. | Housekeeping |
| Tbp | NM_013684. | Housekeeping |
| Tubb5 | NM_011655. | Housekeeping |
